# Revisiting Cucuruchú, a Late Pleistocene site on the Venezuelan Caribbean coast with megafauna and lithic association

**DOI:** 10.64898/2026.06.24.734162

**Authors:** Jorge D. Carrillo-Briceño, Andrés E. Reyes-Céspedes, Arturo Jaimes, Miguel Zabala Reyes, Edwin Alberto Cadena, Rodolfo Sánchez, Alfredo A. Carlini, Antoine Zazzo, Marcelo R. Sánchez-Villagra

## Abstract

The study of historical documents from the original excavation led by J. M. Cruxent, of museum collections, as well as results of fieldwork, are combined to provide a revised assessment of the archeologically site of Cucuruchú, which together with those of Taima-Taima and Muaco, document megafaunal lithic remains associated from the Late Pleistocene in northern Venezuela. Six vertebrate taxa are reported for Cucuruchú: five megaherbivores (a giant sloth, *Eremotherium laurillardi*.; a glyptodontid, *Glyptotherium* cf. *G*. *cylindricum*; a macrauchenid, cf. *Xenorhinotherium bahiense*; a proboscidean, *Notiomastodon platensis*; and an equid, *Equus* sp.), and a turtle (*Kinosternon* sp.). Two fragments of El Jobo projectiles were found *in situ* lying among megaherbivores bones, while another unidentified projectile fragment was also found in the same layer. Although no direct evidence of human exploitation on these megaherbivores remains has been found, the presence of El Jobo projectiles in the undisturbed fossil-bearing layer some of which are associated with fossil remains, could be a suggestive of such interaction. Two new radiocarbon dates indicate ages of at least ca. 15.3-16.0 kyr cal BP for the fossil-bearing layer, contrasting with the mid to late Holocene age originally reported. Cucuruchú is a window to the past and supports the presence of hunter-gatherers of the El Jobo culture in the region at the end of the Pleistocene.

## Introduction

The gateway to South America used by the first humans at the end of the Pleistocene occurred in some points of the vast region that today comprises Colombia and Venezuela, with probable migration routes along the Pacific and Caribbean coasts (Politis *et al*. 2009). In the latter, in the northwestern region of Venezuela, several Late Pleistocene archaeological sites were discovered between the 1950s and 1970s, providing evidence of extinct megaherbivores (body mass > 44 kg) exploitation in association with lithic technologies (Bryan *et al*. 1978; Gruhn and Bryan 1991; Oliver and Alexander 2003; Aguilera 2006; Carrillo-Briceño 2015; Jaimes *et al*. 2024a, b; Carrillo-Briceño *et al*. 2026). These sites are Muaco, Taima-Taima, and Cucuruchú, located in the vicinity of the coastal area comprised between La Vela de Coro and the Taratara town, Falcón State, and separated from each other by only a few kilometres (Figure 1a; Supplementary Information 1). Former radiocarbon dating of the Muaco and Taima-Taima sites, were older than those at sites in North America and the first to contrast with the explanation that was once standard at the time of Clovis-first arrival of people in the Americas (Bryan *et al*. 1978).

**Figure 1.**
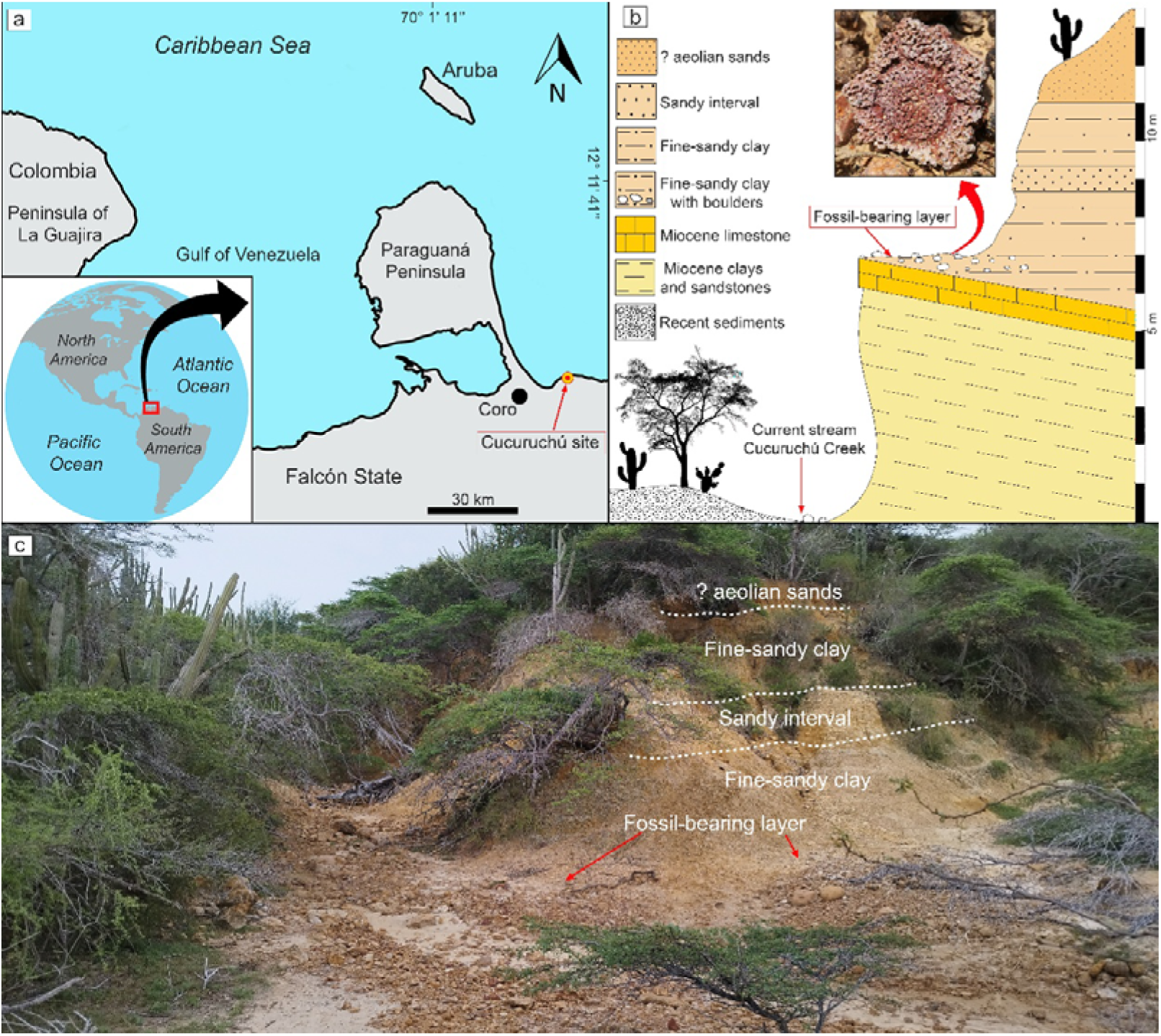
Geographic location of the Cucuruchú site (a), stratigraphic section (b), and overview of the site showing the relative position of the layers (c). Figure “b” shows an osteoderm of Glyptotherium cf. G. cylindricum on the surface of the fossil-bearing layer.

The most representative of these sites is Taima-Taima (Cruxent 1967; Casamiquela 1979; Aguilera 2006; Carrillo-Briceño 2015). There, El Jobo projectile point technology (Cruxent and Rouse 1956; Jaimes *et al*. 2024a; Vargas *et al*. 2025), as well as scrapers, were found in direct association with individuals of the proboscidean *Notiomastodon platensis* (Bryan *et al*. 1978; Cruxent 1978; Ochsenius and Gruhn 1979; Haynes 2022; Carrillo-Briceño *et al*. 2026). Despite the evidence found in Taima-Taima (e.g., Bryan et al., 1978; Ochsenius and Gruhn, 1979), the site has remained controversial and still not universally accepted after the skepticism (without any additional empirical analysis) argued by Lynch (1990) about the lacked solid pre-Clovis human evidence in Taima-Taima and other early South American sites. Lynch’s skepticism and arguments were challenged by Gruhn and Bryan (1991), who clarified several details of previous work and context from Taima-Taima. In addition to the previous evidence presented for Taima-Taima, evidence of probable slaughtering activities in bones of mylodontids (*Glossotherium*), macrauchenids (cf. *Xenorhinotherium*), a toxodontid (?*Mixotoxodon*) (Carrillo-Briceño *et al*. 2026), and glyptodonts (*Glyptotherium* cf. *G*. *cylindricum*) (Carlini *et al*. 2022) have also been reported. Radiocarbon dates for the fossiliferous strata of Taima-Taima suggest an age of 15.3–17.8 cal kyr BP (Carrillo-Briceño *et al*. 2026).

The Muaco and Cucuruchú sites were excavated in 1959 and 1969, respectively, and at both abundant faunal remains and El Jobo projectile points were also found (Royo y Gómez 1960; Rouse and Cruxent 1963; Cruxent 1970; Bocquentin-Villanueva 1979; Oliver and Alexander 2003; Carrillo-Briceño 2015). Of the two El Jobo lanceolate points collected at Muaco, the one in fragmented condition was found *in situ* during the excavation and mixed with the remains of megaherbivores, while the other, in complete condition, was collected on the surface of the deposit (Rouse and Cruxent 1963). The oldest dating of Muaco based on bones (Cruxent 1961) yielded an age of 16,375 ± 400 yr BP, but unfortunately, the presence of prehistoric and modern pottery and glass in the top layer of this deposit (Bryan 1973) generated a debate about the credibility of the association between humans and megafauna at the site. Nevertheless, a bone fragment likely belonging to an extinct megaherbivore from Muaco (see Oliver and Alexander 2003, fig. 17; Aguilera 2006, pp. 28; Carrillo-Briceño 2015, fig. 151), shows a set of at least nine linear, deep, and parallel marks, which could be more associated with an anthropogenic modification than one produced naturally (for example, trampling or marks produced by scavengers).

In the fossiliferous strata of Cucuruchú, which showed no disturbance of any kind (as the top layer of Muaco deposit), two fragments of El Jobo projectiles were found *in situ* lying among megaherbivores bones, while another projectile fragment was also found in the same layer (Cruxent 1970; Bryan 1973). Radiocarbon dating on fossil bones from Cucuruchú gave ages of 3350 ± 160 BP and 5860 ± 80 BP (Tamers 1969), suggesting a mid to Late Holocene age (see Table 1). It is noteworthy that these dates were obtained prior to the setup of the first proper collagen dating protocols (Longin 1971), suggesting that contamination cannot be excluded.

**Table 1.** Chronometric data for the Cucuruchú site. New radiocarbon dated tooth enamel samples are from year 2025. Previous dates are taken from Tamers (1969). Abbreviations: cal yr BP, calibrated years before Present [calibration using the IntCal20 curve (Reimer et al., 2020)]; F-bl, fossil-bearing layer.

| Sample /catalog N° | Stratum | Taxonomy/sample | Anatomical element | Year of analysis | Laboratory N° | Date (BP) | ± | cal yr BP (95.4% CI) |
| --- | --- | --- | --- | --- | --- | --- | --- | --- |
| Cucuruchú 103* | ? | Organic carbon | --- | 1969 | IVIC-514-A | 2080 | 90 | 2310 to 1829 |
| Cucuruchú 103* | F-bl | Indet. turtle | Shell fragment | 1969 | IVIC-514-B | 3350 | 180 | 4090 to 3166 |
| Cucuruchú 104* | F-bl | Glyptodontid | Bone | 1969 | IVIC-512-B | 3980 | 70 | 4797 to 4184 |
| Cucuruchú 108* | F-bl | Megatherid (? *Eremotherium*) | Bone | 1969 | IVIC-511-B | 5860 | 80 | 6886 to 6486 |
| MCH-PV-867 | F-bl | cf. *Xenorhinotherium bahiense* | Lower molariform | 2025 | ECHo6431 | 12960 | 40 | 15657 to 15316 |
| MCH-PV-728 | F-bl | *Notiomastodon platensis* | Molariform | 2025 | ECHo7503 | 13210 | 45 | 16018 to 15700 |

Here, we provide a reassessment of the faunal diversity, stratigraphy, depositional environment and an updated chronology of Cucuruchú. This archaeological site, as well as Muaco and Taima-Taima, are critical for tracing the arrival of humans in South America during the Pleistocene–Holocene transition, their interaction with extinct megaherbivores, as well as the technologies used for fauna exploitation in the northern neotropics (Vargas *et al*. 2025).

### Geographical and historical context

The Cucuruchú site (11° 30′ 10′′ N, 69° 30′ 17′′ W; ∼ 20 m above sea level) is located about 21 km northeast of the Coro city (Falcón State) (Figure 1a), two km north of the Taratara town, and about 1.9 km northeast of the Taima-Taima site (electronic supplementary material S1). The site is located on the right bank of the intermittent Cucuruchú Creek, near its opening to the Caribbean Sea, about 180 m upstream the shoreline. The sediments containing the fossiliferous layer of the archaeological site rest unconformably on Miocene-age rocks, a section that was exposed due to erosion/weathering at the Cucuruchú Creek (Figure 1b). A small, superficially dry gully erodes in an east-west direction and descends to the bottom of the Cucuruchú Creek, where it gives rise to a small spring known as Agua Divina. The current vegetation around the Cucuruchú is dominated by xerophytic shrubland (Figure 1c), characteristic of most of the northern Falcón State (Ochsenius 1980).

Although Cruxent (1970) mentioned that the Cucuruchú site was excavated during April 1969 (Figures 2 and 3), his site’s field notebook suggests that the excavation began on March 21 of the same year. There is no reliable information on how long the excavation at Cucuruchú lasted, but probably from the end of March until May 1969, following Cruxent’s (1970) statement that in June of the same year they resumed excavations at the Taima-Taima site. The excavation encompassed a rectangular area of 14 x 8 m divided into 28 sections, each 2 m on a side (see Cruxent 1970), defined in three main areas designated by Cruxent in his field notebook as CX-104, −105, and −108 (electronic supplementary material S1). According to Cruxent (1970), the fossil-bearing layer (see Figures 1b, 2 and 3) was dry, with undisturbed stratigraphy, thicknesses between 40 cm and 100 cm, and characterized by sediments with a clay matrix and abundant rounded limestone fragments ranging from pebbles to boulders.

**Figure 2.**
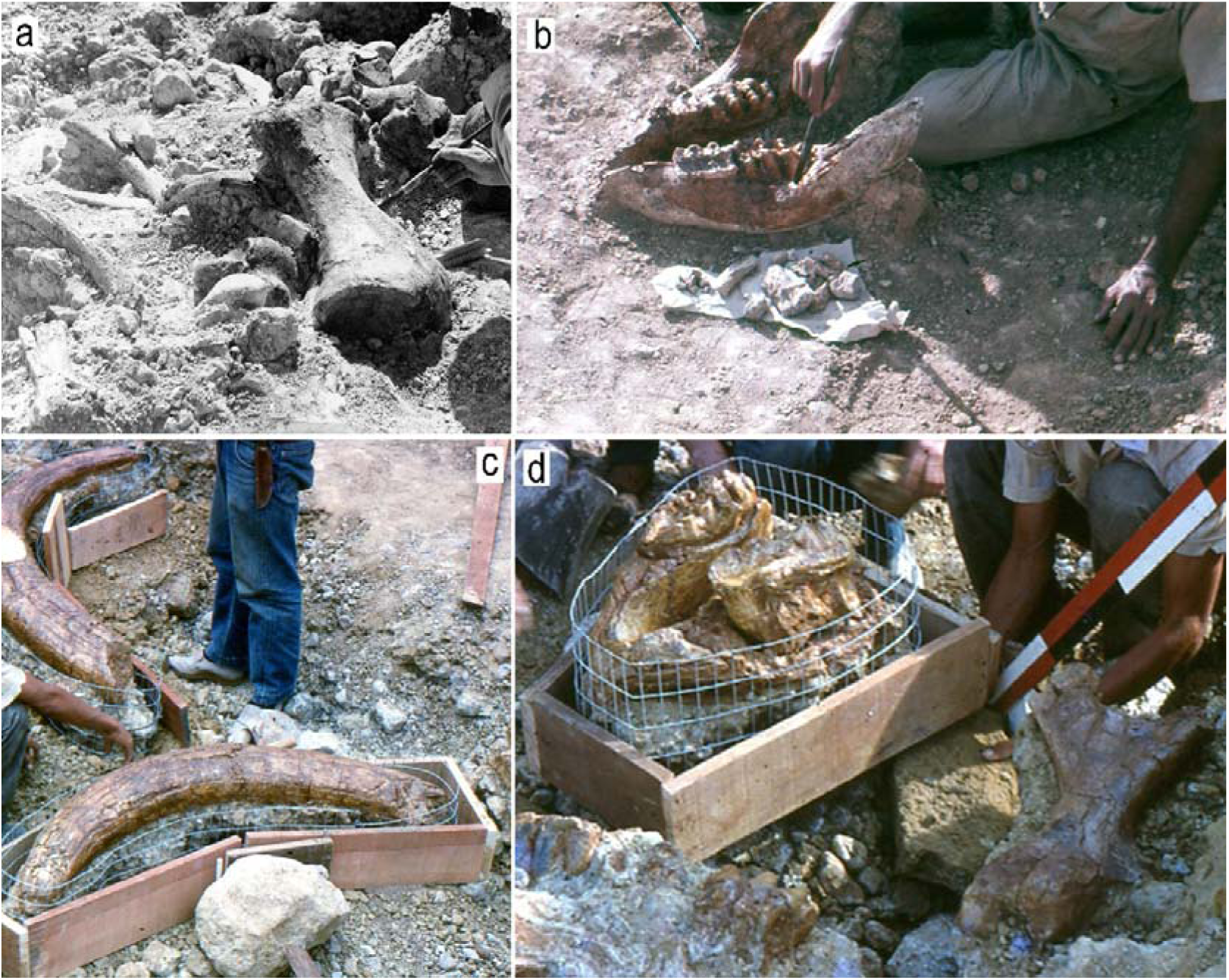
The Cucuruchú site during the excavation of 1969. Carcass of *E*. *laurillardi*. (a) showing a left tibia (see Figure 4a–d), astragalus, vertebrae and ribs. Complete mandible (b) of a probable subadult or adult of *Notiomastodon platensis* excavated in grid CX-108. Tusks, mandible, a fragmented maxilla with upper molars (c, d) of an adult of *Notiomastodon platensis* excavated in grid CX-105. In the lower right corner of figure “d” a left femur likely belonging to *Glyptotherium* can be seen. Images courtesy archive of the Universidad Experimental Francisco de Miranda (a), Fundación José María Cruxent in the custody of the Laboratorio de Archeología of the Instituto Venezolano de Investigaciones Científicas (b–c).

**Figure 3.**
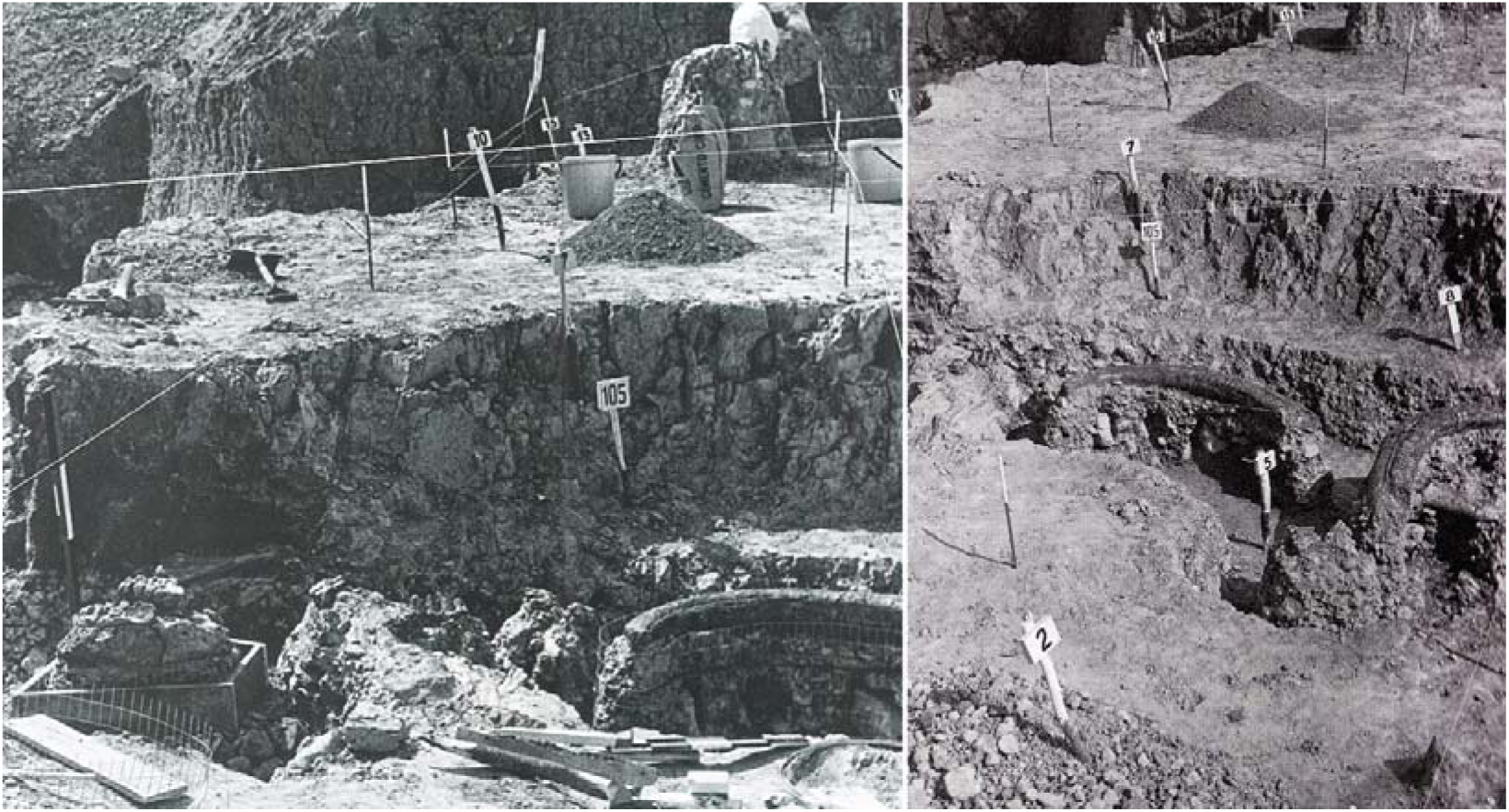
Remains of an adult of *Notiomastodon platensis* excavated in area CX-105 (remains also shown in Figure 2). These two images were erroneously identified in Cabrero 2009 as belonging to the Taima-Taima site. Modified images of Cabrero (2009), with permission of Instituto Venezolano de Investigaciones Científicas.

A preliminary list of the fossil taxa of Cucuruchú was made by the French paleontologist Robert Hoffsteter, and the reported assemblage included “*Haplomastodon guayanensis*, *Eremotherium rusconii* and *Glyptodon clavipes*” (Cruxent 1970). No subsequent study of the taxonomic and taphonomic aspects of the fossil assemblage was ever published, and it is unknown whether any catalogue of the assemblage ever existed. To our knowledge, all the collected specimens were deposited at the Instituto Venezolano de Investigaciones Científicas (IVIC), and some of these specimens are still identified with the section of the excavation from which they were collected (e.g., CU-F-108, see Figure 4n–p). In reference to the three fragments of El Jobo-type lithic projectiles (Cruxent 1970; fig. 3; Bryan 1973), two of them were found lying among the fossil bones remains (Figure 5; electronic supplementary material S1), and the third in the same layer but without association to fossils.

**Figure 4.**
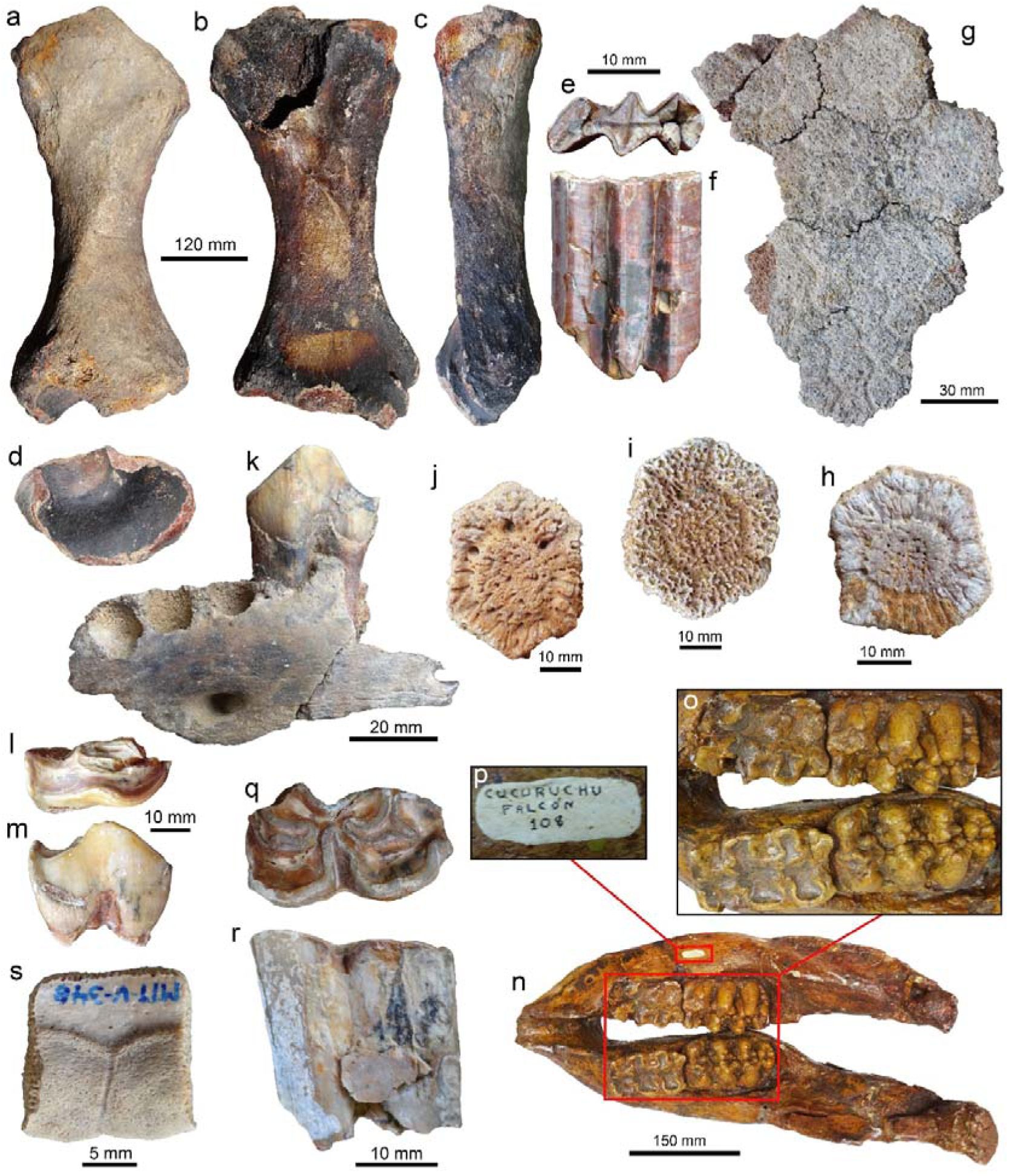
Fossil remains from the Cucuruchú site. Left tibia (a–d: IVIC-AP-023) of *Eremotherium laurillardi* with burned surfaces probably because of the 1978 fire at IVIC. Upper left molar (e, f: MTT-V-297), fragment of armoured carapace (g: MCH-Pv-870), and isolated carapace osteoderms (h: MCH-Pv-912; i: MCH-Pv-903; j: MCH-Pv-916) of *Glyptotherium* cf. *G*. *cylindricum*. Fragment of the left hemimandible with premolar 3 (k) and isolated premolar (?p4) (i, m) (MCH-Pv-870) of cf. *Xenorhinotherium bahiense*. Mandible (n–p: IVIC-Cu-F-108) of a subadult of *Notiomastodon platensis*. Isolated lower left premolar p3 or p4 (q, r: MTT-V-323) of *Equus* sp. Peripheral plate (s: MTT-V-348) of the turtle *Kinosternon*. Specimens a–d and n–p come from the 1969 excavation. Views: anterior (a), dorsal (m), posterior (b), distal (d), external (g–j, s), labial (m), lateral (c, k), lingual (r), occlusal (e, I, o, q), indet. (f).

**Figure 5.**
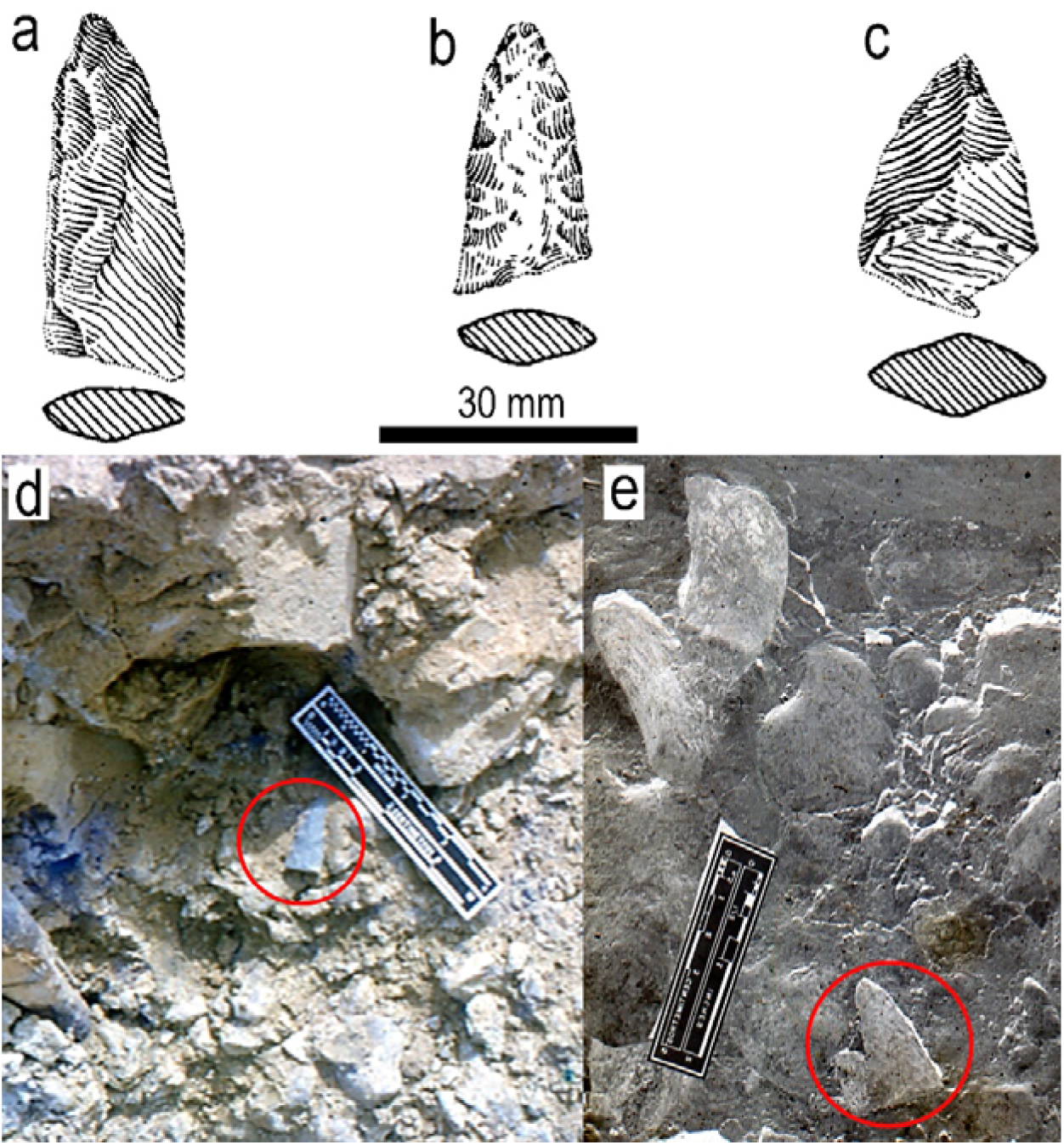
El Jobo projectile fragments found at the Cucuruchú site (**a**–**e**). Drawing (including view of the section) of the three projectile fragments (**a**–**c**: modified from Cruxent 1970). According to Cruxent (1970), projectiles **a** and **b** were found among fossil bones. Projectiles in situ during the excavation of 1969. (**d**, **e**). Images (**d**, **e**) courtesy archive of the Fundación José María Cruxent and Instituto Venezolano de Investigaciones Científicas.

## Materials and methods

### Stratigraphic reassessment

A new stratigraphic survey of the section containing the Cucuruchú site was carried out by the authors (A.E.R.C.; J.D.C.B.; R.S.) in May of 2025. The sediments corresponding to the fossil-bearing layer of the archaeological site rest unconformably on an outcrop of Miocene-age (Caujarao Formation) that has been vertically cut by erosion. The section (Figure 1b) was measured from the base at the current stream of the Cucuruchú Creek to the top, the latter represented by the sediments overlying the fossil-bearing layer of the archaeological site. Additional information on sedimentology, stratigraphy, and excavation context was obtained from the data compiled by J.M. Cruxent in his site’s field notebook, which had not been published, from the historical archive of the archaeology laboratory of the Instituto Venezolano de Investigaciones Científicas (IVIC), Miranda State.

### Taxonomy

We studied the collection of fossil vertebrates excavated at the Cucuruchú site by Cruxent and team in 1969 (Cruxent 1970), as well as other new fossil specimens’ we collected at the site during the last ten years. Some of the specimens collected in the 1969 excavation are deposited in the collection of the archaeology laboratory of the IVIC, while others were probably lost in the fire that affected the deposits of same institution in 1978 (e.g., Figure 4a–d). Cruxent (1970) did not specify the total number of fossil specimens or an approximate number of individuals per taxon collected during the 1969 excavation in his publication or in his site’s field notebook. He also did not specify which bone materials were in association or assign any catalog numbers to them; although some specimens preserve the provenance number of the excavation grid where they were collected (e.g., Figure 4p). Thanks to the use of historical photographic record of the excavation and the data from the site’s field notebook (electronic supplementary material S1), we have been able to confirm that some of the bone materials deposited in the IVIC collection (without a prior provenance record associated) come from the Cucuruchú site.

In reference to the new fossil specimens reported herein (n= 42), those were collected by several of the authors during rescue paleontology activities on several occasions between 2015 and 2025. The specimens were surface-collected and come from the same fossil-bearing layer excavated in 1969 (see Figure 4). The new specimens are housed in the Museo Comunitario de Taratara “Cristóbal Higuera” (MCH-Pv- and MTT-V), Falcón State. Other specimens from the Cucuruchú site includes a partial end of the caudal tube (CIAAP-1540) of a glyptodontid (see Carlini *et al*. 2008, fig. 3E, F), probably surface collected in the 1980s–90s and deposited in the Centro de Investigaciones Antropológicas, Arqueológicas y Paleontológicas (CIAAP) de la Universidad Nacional Experimental Francisco de Miranda (UNEFM), Coro. The taxonomic identification provided here involved an extensive bibliographic review and comparisons with fossils specimens. National collections studied for comparative purposes that preserve Late Pleistocene fossils from Muaco, Taima-Taima, and other localities of Falcon State and Venezuela, included: UNEFM-CIAAP, IVIC; Museo de Ciencias de Caracas (MCNC); Museo Geológico Royo y Gómez de la Universidad Central de Venezuela (UCV), Caracas; y Fundación La Salle de Ciencias Naturales, San Carlos, Cojedes State.

### Radiocarbon dating

Tooth enamel dating was performed on two fossil teeth from the Cucuruchú site at the radiocarbon laboratory of the Muséum national d’Histoire naturelle (MNHN), Paris and AMS 14C dating was performed using the AMS ECHoMicadas system at LSCE (Saclay). The dated tooth enamel comes from a premolar (left p4M, CH-PV-867) of the macrauchenid cf. *Xenorhinotherium bahiense*, and a molar fragment of determined position of *Equus* sp. (MCH-PV-728), with their stratigraphic provenance shown in Figure 1b. The enamel surface of the specimens was cleaned of sediments, and the dentine was removed with a Dremel to isolate the enamel from the rest of the dental tissue. Then, the enamel (∼2 g) was grounded using a steel mortar and pestle, followed by grinding in an agate mortar to a particle size of < 100 microns. The powder was then finely (5-10 microns) crushed using a McCrone Microniser Retsch following (Wood *et al*. 2016), as this approach increases the efficiency of the acetic acid pre-treatment aimed at removing diagenetic carbonates and therefore improves the radiocarbon ages obtained. The resulting powder was pre-treated under a light vacuum for 20 hours with a solution of 1 N acetic acid then rinsed with milliQ water and oven-dried at 50°C. About 250 mg of powder was then reacted under vacuum with orthophosphoric acid at 70 °C for around 20 min. The CO2 gas produced was then reduced in the presence of hydrogen and iron to produce graphite. Samples were then pressed into targets and ^14^C ages were measured on the compact AMS ECHoMicadas at LSCE, Saclay.

All the samples were collected and transported for study with authorization from the Instituto del Patrimonio Cultural de Venezuela (IPC) using permissions: VE-IPC-CEBC-PP-06/2022-1 and VE-IPC-CEBC-PP-01/2023.

## Results

### Stratigraphy and depositional environment

The stratigraphic section, measured from the base of the current stream of Cucuruchú Creek to the top, has an approximate thickness of 14 meters (Figure 1b). The first seven meters correspond to outcrops of clays, sandstones, and limestones of the Mataruca Member of the Caujarao Formation (Late Miocene). The top of this Miocene unit in the section is characterized by a fossiliferous limestone, overlain by Quaternary sediments referred to here as the fossil-bearing layer of the Cucuruchú site. The sediments of the fossil-bearing (Figure 1b, c) layer are horizontally positioned, with a thickness up to ∼100 cm. This non-heterogeneous and poorly selected layer is characterized by an unconsolidated matrix of fine-sandy clay with abundant pebbles and boulders of limestone and sandstone, ranging from angular to rounded of different sizes and up to a little over 15 cm of diameter. Cruxent (1970) and Cruxent and Ochsenius (1979) considered that the accumulation of bones together with boulders and rounded pebbles in this layer was due to alluvial deposition in the talweg of an intermittent watercourse. However, given the characteristics of the deposit, a colluvial origin is more likely (see discussion below). The El Jobo projectiles reported by Cruxent (1970) come from this layer. Almost all the new fossil specimens reported here (except for the articulated osteoderms MCH-Pv-870, see below) were collected on the surface, still embedded in the matrix, from the section of the layer exposed during excavation of 1969, affected since then by erosion.

Overlying the fossil-bearing layer there is a greenish-gray fine-sandy clay layer that weathers in color yellow ochre. This unconsolidated layer contains horizontally deposited sediments and reaches a thickness of approximately two meters. We collected some glyptodontid osteoderms (e.g., MCH-Pv-870; Figure 4g; Supplementary Information 1) at the base of this layer, a finding consistent with similar observations made by Cruxent (1970); so far, this is the only sediments, apart from the fossil-bearing layer, in which the presence of fossil vertebrates could be referenced at the site. Above the later layer, there is a sandy interval, ranging in color from brown to ochre, with an average thickness of approximately 40 cm. This sandy interval is followed by a greenish-gray fine-sandy clay accumulation that weathers in color to yellow ochre. This layer is horizontal and unconsolidated, with a thickness of approximately 2 m. The fine size of these sediments (as well as the ones overlaying the fossil bearing layer), their good selection and position of horizontal layers, suggest that they must have been deposited in a low-energy, protected or restricted environment, probably a lagoon. All the layers mentioned above were deposited long before the Cucuruchú Creek cut through the site.

The top of the section (Figure 1c) is characterized by unconsolidated reddish-ochre sands of probable aeolian origin. The current soil supporting the vegetation cover of the area forms the top of this sand.

### Faunistic assemblage

Six vertebrate taxa are reported here for the fossil-bearing layer of the Cucuruchú site. The mammals are megaherbivores: the terrestrial sloth *E*. *laurillardi* (Figures 2a, 4a–d), the glyptodontid *Glyptotherium* cf. *G*. *cylindricum* (Figure 4e–j), the macrauchenid cf. *Xenorhinotherium bahiense* (Figure 4k–m), the proboscidean *Notiomastodon platensis* (Figures 2b–d, 4n–p), and the equid *Equus* sp. A pond turtle belongs to the genus *Kinosternon* (Figure 4s). The taxa cf. *Xenorhinotherium bahiense*, *Equus* sp., and the *Kinosternon* sp., are reported for the first time for the Cucuruchú site; previously indeterminate turtle remains had been mentioned for the site (Tamers 1969). Taxonomic assignment and other remarks on the faunal assemblage are documented in the electronic supplementary material S1.

At least six individuals of megaherbivores are reported here for the Cucuruchú site. A conservative estimate of the minimum number of individuals per species is one (except for gomphotheres, see below); a minimal number as many of the specimens collected in 1969 were found disarticulated at the site (e.g., *Glyptotherium* cf. *G*. *cylindricum*), it difficult to associate them with a particular individual. We recognize at least one individual, probably an adult, of *E*. *laurillardi*, corresponding to the remains excavated in 1969 (Figure 2a). Two individuals of *N*. *platensis* are identified, including the mandible, a fragmented maxilla with the upper molars and two long tusks of an adult excavated in area CX-105 (see Figures 2c, d and Figure 3; Supplementary Information 1), and the complete mandible excavated from CX-108, probably belonging to a subadult individual (Figure 2b and 4n). The mandibular remains assigned here to cf. *Xenorhinotherium bahiense* (MCH-Pv-867) likely corresponds to a single individual. The two molars of *Equus* sp. (MCH-Pv-728 and MTT-V-323) may or may not belong to the same individual. The remains assigned to *Glyptotherium* cf. *G*. *cylindricum* are the most abundant at the Cucuruchú site, and these include remains of the carapace, fragments of the end of the caudal tube (Supplementary Information 1) and fused tail rings, abundant isolated osteoderms and a single molariform (e.g., Figure 4e–j). The assignment of these isolated materials to one or more individuals is not possible (Supplementary Information 1).

### Age of the Site

The two tooth enamel samples from molars of cf. *Xenorhinotherium bahiense* and *Equus* sp., coming from the fossil bearing layer were dated to 12,960 ± 40 and 13,210 ± 45 BP, which, once calibrated, indicate an age comprised between 15,3 and 16 kyr cal BP (Table 1). This is likely to be a minimum age given the material’s propensity for modern carbon uptake during the fossilization process (Zazzo and Saliège 2011; Zazzo 2014). The sediments overlying the fossil-bearing layer could not be dated due to a lack of fossils from these layers.

### Ancient palaeoenvironment, fauna and cultural evidence in Cucuruchú

The Quaternary deposits that characterize the Cucuruchú site likely have an origin that we associated with two distinct events, the first producing the fossil-bearing layer, and the second depositing the overlying sediments, the latter with the presence of some fossils only at its base (e.g., MCH-Pv-870; Figure 4g; Supplementary Information 1). The first depositional event occurred on a paleorelief characterized by outcrops of the first limestone of the Late Miocene Mataruca Member. The fossil-bearing layer was deposited on top of this limestone, and its heterogeneous and poorly sorted sediments, with abundant pebbles and boulders, suggest a colluvial origin. Cruxent (1970) suggested that, due to the roundness of the pebbles and boulders, they originated several kilometers south of the Cucuruchú site. Notwithstanding, our interpretation suggests that a primary source where sediments containing the pebbles and boulders, upon which vertebrate remains accumulated, were close to the final place of deposition, and at some point, they were displaced and mixed, probably by mass wasting. There is no detailed report on the taphonomic aspects of the fossils excavated at the Cucuruchú site in 1969. However, the photographic record of the excavation supports our hypothesis regarding a nearby source of origin for the skeletal remains before their final deposition in the current fossil-bearing layer. Some of the individuals excavated in 1969 were found in a state of semi-articulation or association. For example, an adult *N*. *platensis* individual excavated in area CX-105 (Figures 2c, d and 3; Supplementary Information 1) has a mandible, a fragmented maxilla with the upper molars *in situ*, two long tusks, and what appears to be a long bone, all found in association. Similarly, the *E*. *laurillardi* individual (Figure 2a) has abundant associated bone elements. The bone materials that we have been able to analyze from the IVIC collection, and the new ones collected by us do not show evidence of erosion by transport. The fragmentation of some specimens (e.g., the mandible cf. *Xenorhinotherium bahiense*) likely occurred after they were exposed to the surface on the site and affected by weathering and other erosive agents for years.

The cause of the accumulation of carcasses and other isolated elements in the primary source deposit and subsequently transported, mixed and embedded in the fossil-bearing layer, is unknown. This single or multiple accumulation event of megaherbivore carcases and other animal remains in the primary source deposit could be related to the natural death or the result of predation by carnivores, although the exploitation and use of these remains as resources by humans should not be ruled out, mainly considering that seems to be a selectivity to big animals (e.g., Ben-Dor and Barkai 2024)

During the 1969 excavation, three lithic artifacts embedded in the fossil-bearing layer *in situ* were found in Cucuruchú, identified as fragments of El Jobo-type projectiles (Cruxent 1970; Bryan 1973). According to Cruxent (1970, fig. 3), two of these were found in situ lying among megaherbivores bones (see Figure 5d, e; Supplementary Information 1), while another projectile fragment (Figure 5c) was also found in the same layer but not associated with fossils. According to Bryan (1973), one of the projectiles was found adjacent to a proboscidean bone. These projectiles could not be located in the IVIC collection; however, based on the illustrations presented by Cruxent (1970), we believe that two of them could correspond to fragments of the proximal section of the projectile body (Cruxent 1970, fig. 3.1, 2) and are likely similar to morphotype 3 of this lithic technology (Vargas *et al*. 2025). The third fragment (Cruxent 1970, fig. 3.4) was not associated with the fossil remains, and we hypothesize it is a projectile preform. There is no evidence to suggest that these lithic artifacts were in direct physical association with the carcasses in the primary source deposit. The megaherbivores remains from the 1969 excavation we examined show no evidence of potential anthropogenic modification. However, it is highly probable that these lithic artifacts lay in the same primary source sediments where the megaherbivore carcasses accumulated, subsequently becoming part of the fossil-bearing layer. Bryan (1973) even noticed that in Cucuruchú “bones and artifacts have apparently been moved from their original position and redeposited in the gravel, the lack of evidence of water wear on either the bones or the artifacts suggests that their original juxtaposition was very close”. As Cruxent (1970) reported, the fossil-bearing layer was undisturbed after its deposition, remaining completely protected and sealed by the overlying sediments from further disturbance.

The radiocarbon dates of the fossil-bearing layer (Table 1) provide a chronological context of these El Jobo projectiles. The age of the Cucuruchú fossil-bearing layer is contemporaneous with the El Jobo projectile-bearing layers at the Taima-Taima site (Carrillo-Briceño *et al*. 2026). Other Venezuelan sites where El Jobo projectiles have been found in sediments alongside megafauna remains include Muaco (Royo and Gómez 1960; Rouse and Cruxent 1963), and El Vano (Jaimes *et al*. 2024b). In the latter, an *E*. *laurillardi* carcass was found in association with several projectiles (Jaimes *et al*. 2024).

All the megaherbivore taxa referred to here for the Cucuruchú site have also been reported at Muaco and Taima-Taima, and other contemporaneous Late Pleistocene sites in Falcón State (Carrillo-Briceño *et al*. 2024, 2026, Wilson *et al*. 2024, and references therein). These megaherbivores and other animals likely congregated in the ancient environment of Cucuruchú to obtain food or water. Supporting this idea is the fact that *Kinosternon* turtles prefer aquatic environments, inhabiting streams, rivers, lagoons, seasonal or perennial ponds (Iverson *et al*. 2013).

A watering hole likely existed at the ancient Cucuruchú site, influenced by the action of resurgent springs, such as those reported for Muaco and Taima-Taima (e.g., Royo and Gómez 1960; Ochsenius 1980; Gruhn and Bryan 1984; Carrillo-Briceño *et al*. 2026), and other potential sites recently discovered (and under study) by our team in the surrounding area. At Cucuruchú, water emanations known as the Agua Divina spring still exist, originating in the joints of the limestone layer and draining into the current Cucuruchú Creek stream. These water emanations could have been present in the locality since the Late Pleistocene, and this could support our hypothesis of what we refer to here as a second depositional event, represented by the sediments overlying the fossil-bearing layer, which were likely deposited in a low-energy or restricted environment. Cruxent (1970) mentioned the presence of stem and leaf impressions in these sediments, and although we could not find any plant remains in our sampling, these deposits probably represent potential for paleobotanical and palynological studies.

### Final remarks

The remains of megaherbivores and other fossil vertebrates present in the fossil-bearing layer of Cucuruchú are the result of a secondary event. This could have happened due to a mass movement that mixed sediments, clasts, boulders, pebbles, bones, and lithic artifacts, resulting in their transport from the primary accumulation source to the final deposition site in the paleorelief, likely over a short distance between the two.

The Cucuruchú faunal assemblage, represented by five megaherbivore taxa, is like other contemporaneous assemblages from the Late Pleistocene, such as those from Muaco and Taima-Taima (Aguilera 2006; Carrillo-Briceño *et al*. 2026), located just a few kilometers away. The sedimentary characteristics of the Quaternary deposit described here for Cucuruchú, and the existence of a spring emanating from the underlying Miocene rocks, support the hypothesis that this spring has existed at the site at least since the Pleistocene, probably with the presence of a watering hole nearby where animals congregated to drink, as has been inferred for the sites of Taima-Taima (Carrillo-Briceño *et al*. 2026) and Muaco (Royo and Gómez 1960; Carrillo-Briceño 2015).

Although no direct evidence of human exploitation of megaherbivores and other faunal resources has been found at the Cucuruchú site, the presence of El Jobo projectiles in the fossil-bearing layer, some of which are depositionally associated with fossil remains (Cruxent 1970), are suggestive of such interaction. The new dates presented here for the fossil-bearing layer indicates an age of at least 15.3–16.0 kyr BP, which is broadly contemporaneous with the Taima-Taima site (Carrillo-Briceño *et al*. 2026).

The abundance of fossils and the three projectiles, all from a small section excavated at Cucuruchú, suggest that this site still holds potential for further excavations, as it could shed new light on faunal assemblages and technologies used for fauna exploitation in the region at the end of the Pleistocene.

## Supporting information

