## Supplementary material for "Revisiting Cucuruchú, a Late Pleistocene site on the Venezuelan Caribbean coast with megafauna and lithic association"

**ELECTRONIC SUPPLEMENTARY MATERIAL S1.**

**Geographic location and Historical photographic record of the Cucuruchú excavation and data from the site's field notebook**

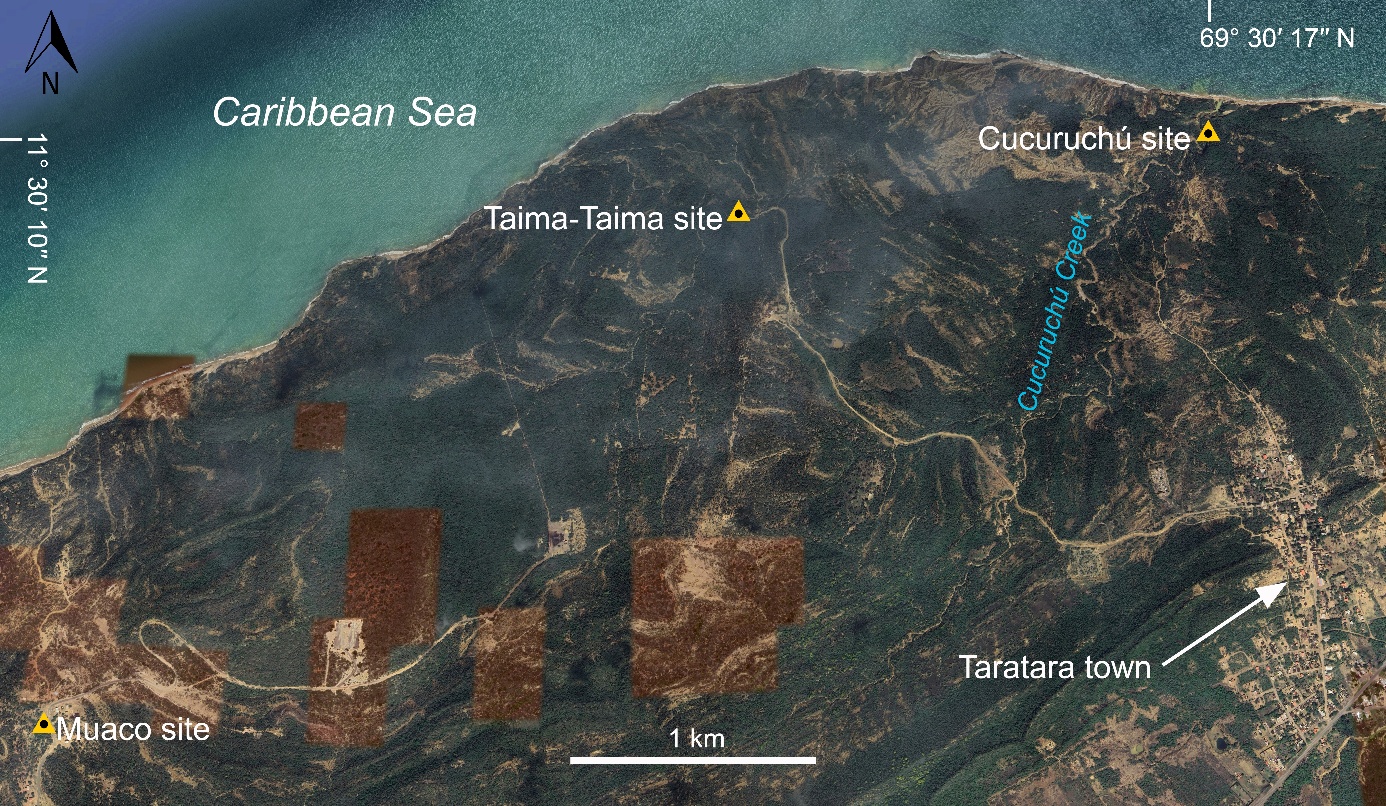

**Figure S1**.**1**. Satellite image showing the location of the Cucuruchú site. Figure based on Google Earth Pro satellite image (2026). https://earth.google.com/ [Accessed March 16, 2026].

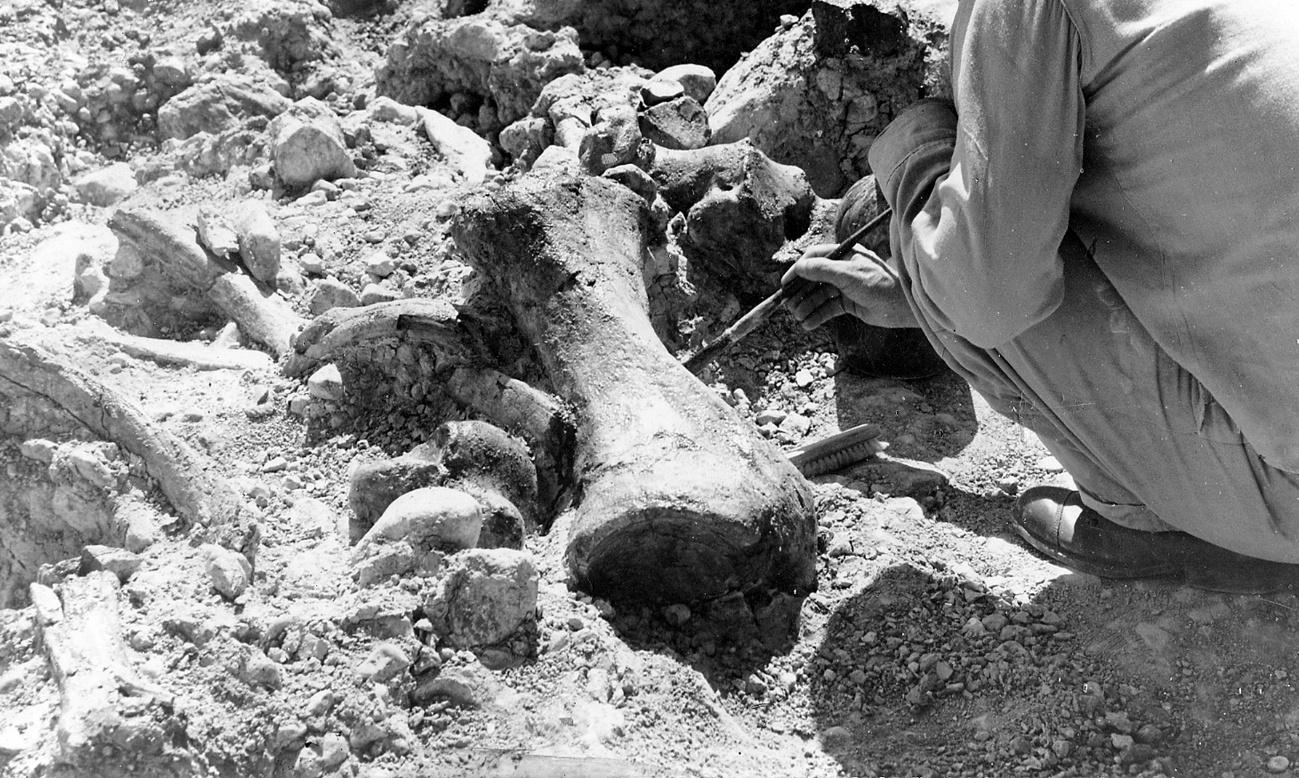

**Figure S1**.**2**. Carcass of *Eremotherium* *laurillardi* at the Cucuruchú site during the excavation of 1969. The image shows a left tibia, astragalus, vertebrae and ribs. Image courtesy archive of the Universidad Experimental Francisco de Miranda.

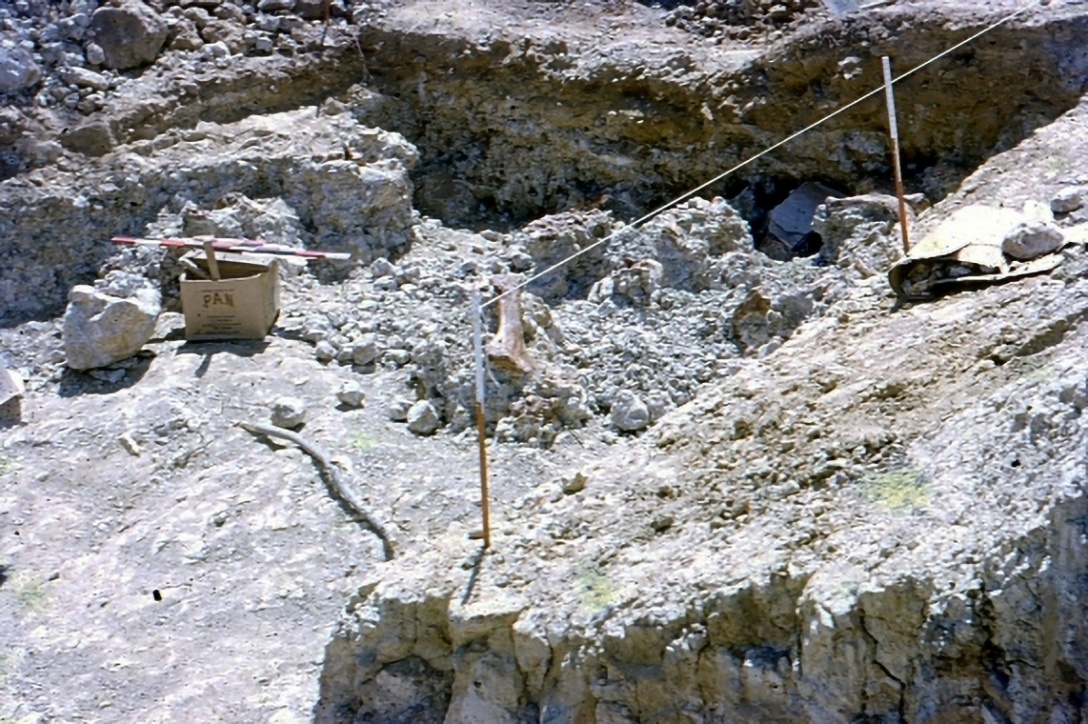

**Figure S1**.**3**. Excavation at the Cucuruchú site in 1969. In the central part of the image, a long-fossil bone of an undetermined mammal can be seen. Imagen from Fundación José María Cruxent in the custody of the Laboratorio de Archeología of the Instituto Venezolano de Investigaciones Científicas.

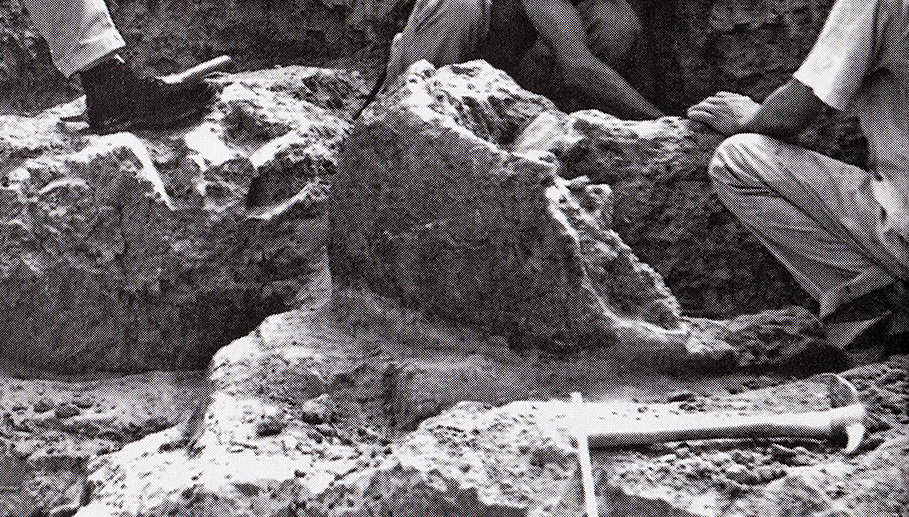

**Figure S1**.**4**. Extraction of the glyptodont carapace fragment in Cucuruchú in 1969. This image is erroneously identified in Cabrero 2009 as belonging to the Taima-Taima site. Modified image of Cabrero (2009), with permission of Instituto Venezolano de Investigaciones Científicas.

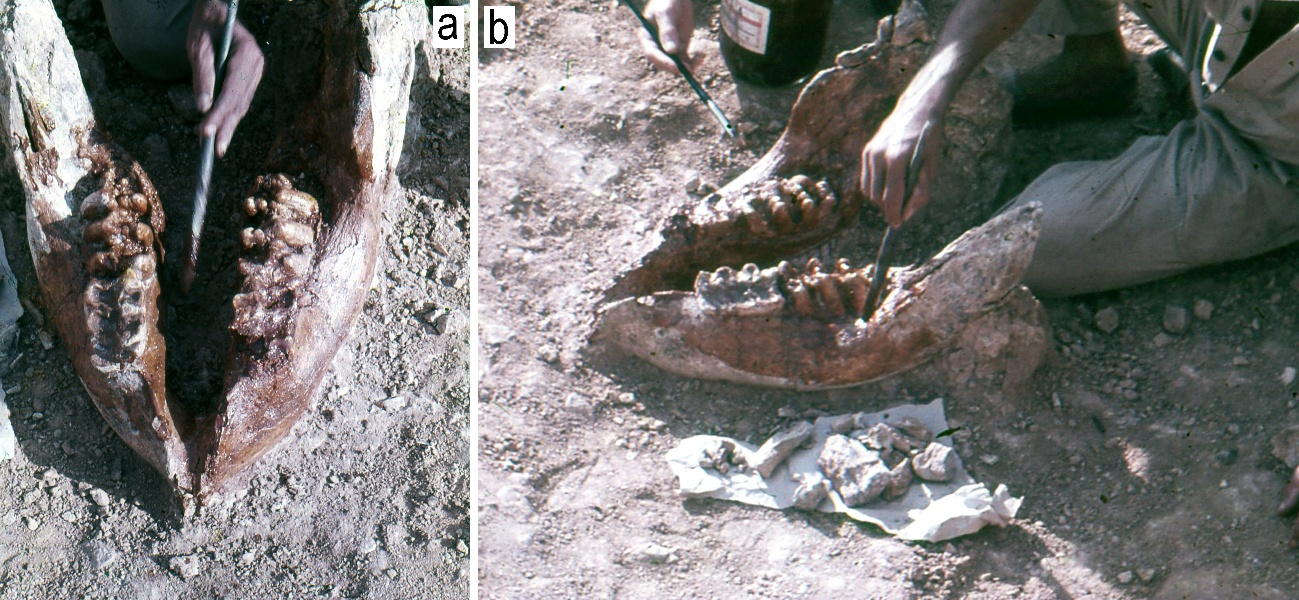

**Figure S1**.**5**. Mandible (**a**, **b**) of a probable subadult or adult of *Notiomastodon platensis* excavated in area CX-108. Image from Fundación José María Cruxent in the custody of the Laboratorio de Archeología of the Instituto Venezolano de Investigaciones Científicas.

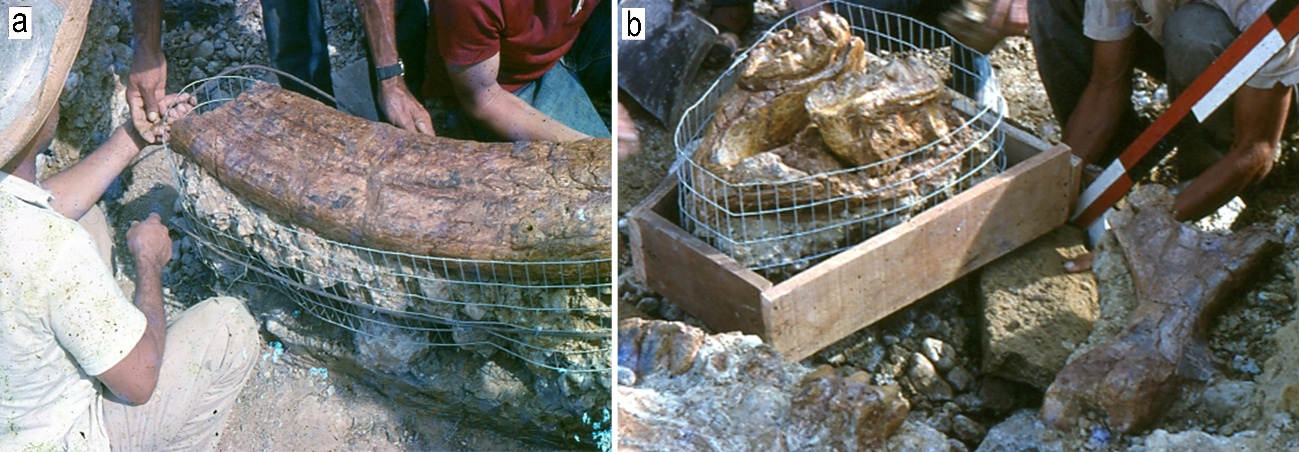

**Figure S1**.**6**. Tusks (**a**), and mandible (b) of an adult of *Notiomastodon platensis* excavated in area CX-105. In the lower right corner of figure **b**, a left femur likely belonging to *Glyptotherium* can be seen. Images from Fundación José María Cruxent in the custody of the Laboratorio de Archeología of the Instituto Venezolano de Investigaciones Científicas.

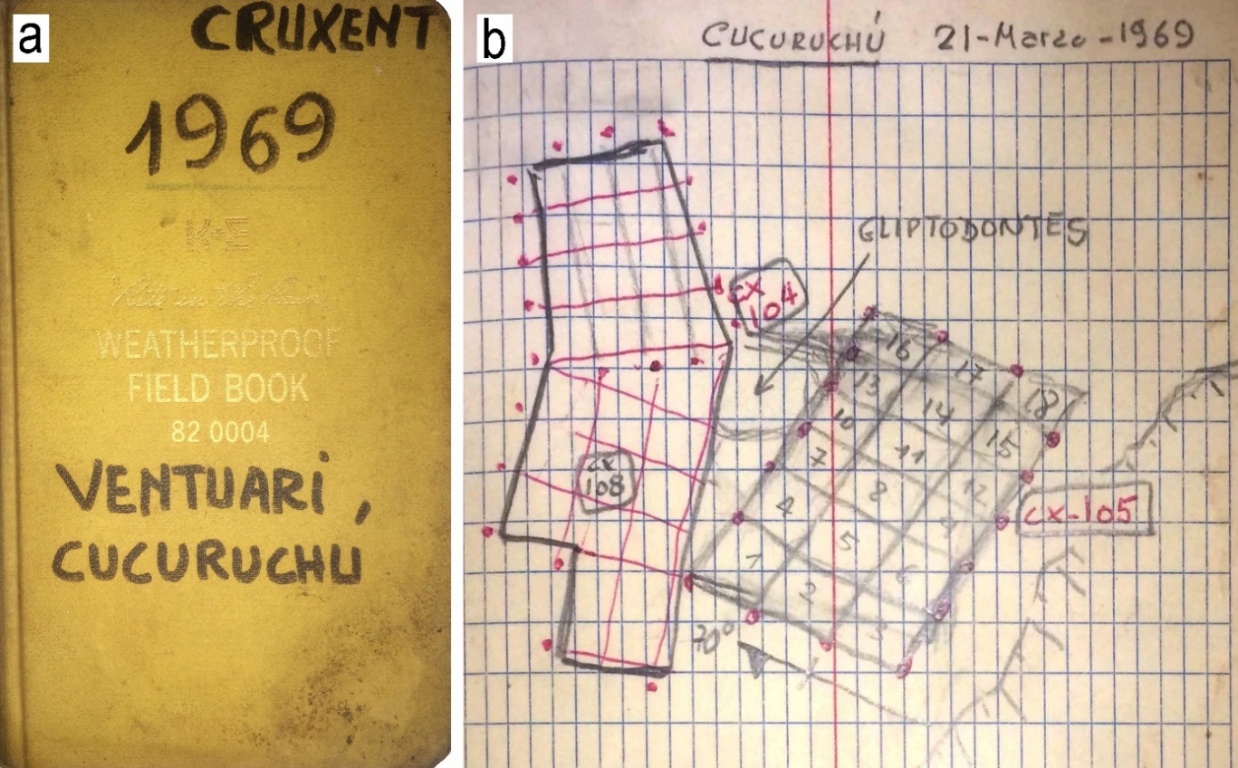

**Figure S1**.**7**. J.M. Cruxent site's field notebook (**a**) with details of the 1969 excavation diagram (**b**). Courtesy Laboratorio de Archeología of the Instituto Venezolano de Investigaciones Científicas.

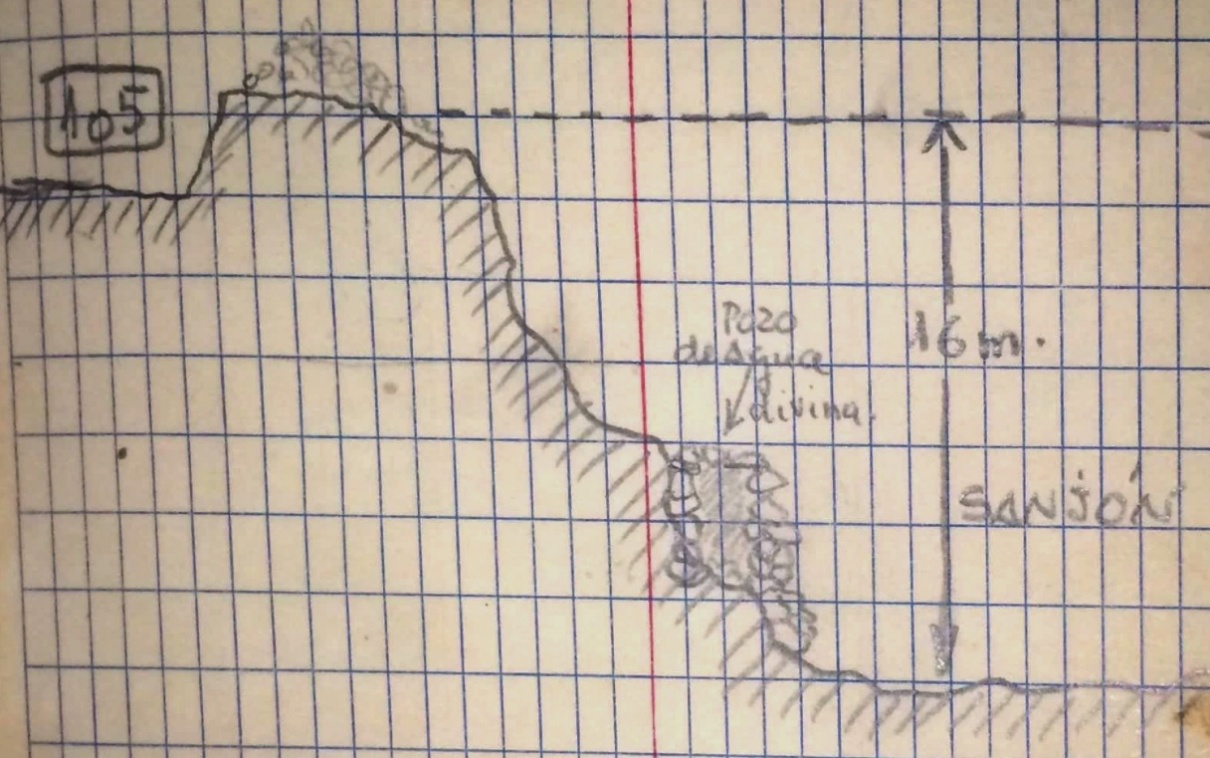

**Figure S1**.**8**. J.M. Cruxent site's field notebook (showing the location of the site in reference to the Cucuruchú Creek. Courtesy Laboratorio de Archeología of the Instituto Venezolano de Investigaciones Científicas.

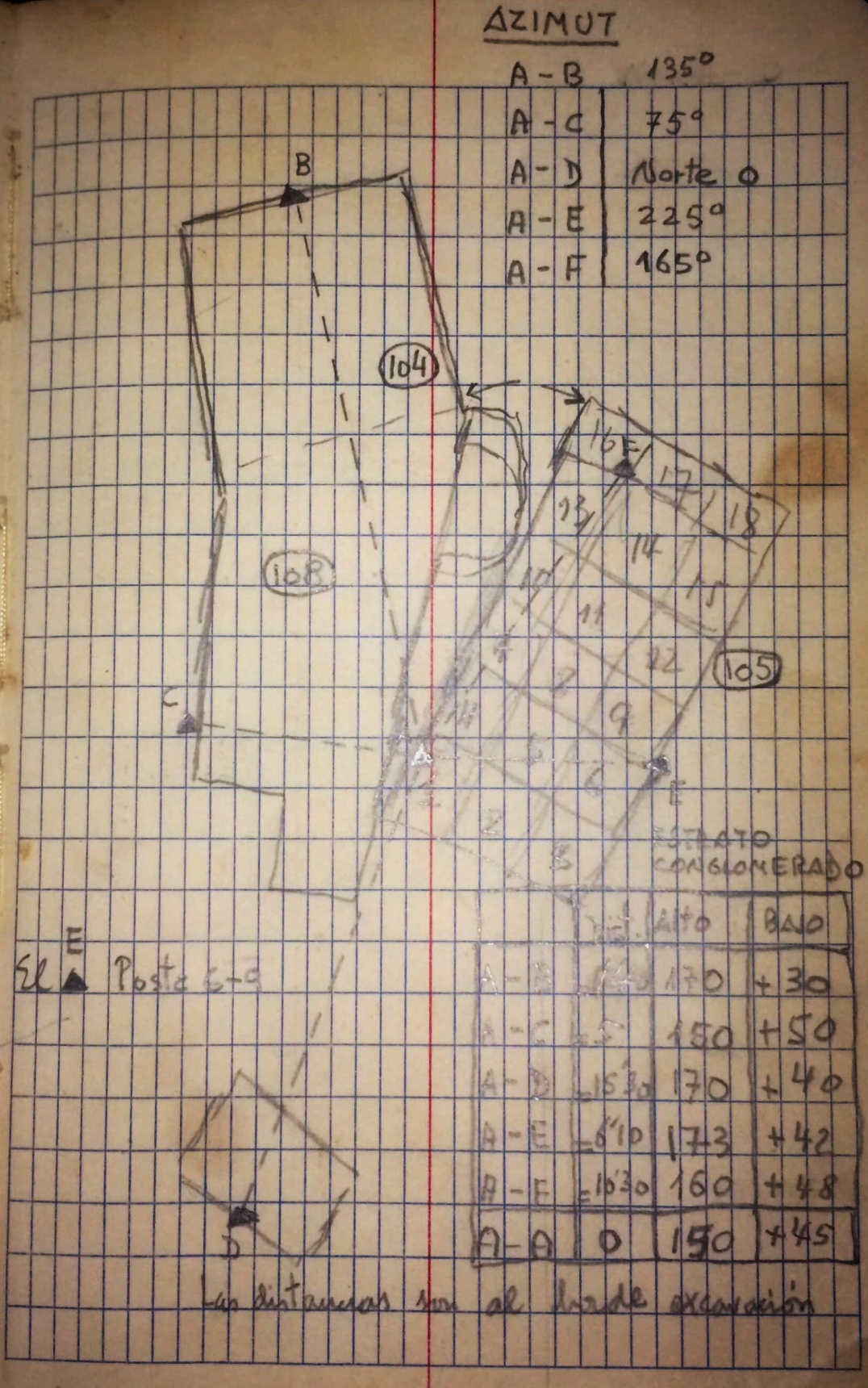

**Figure S1**.**9**. J.M. Cruxent site's field notebook with details of the 1969 excavation diagram. Courtesy Laboratorio de Archeología of the Instituto Venezolano de Investigaciones Científicas.

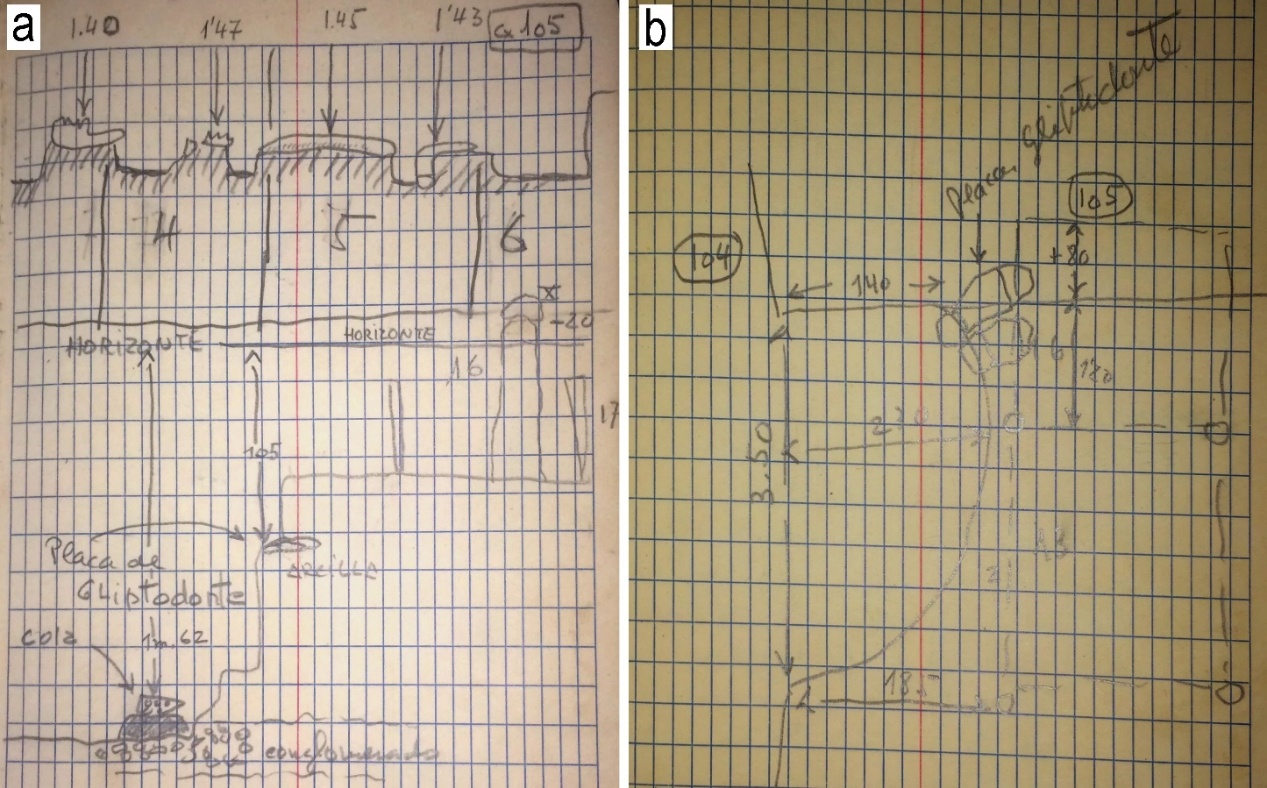

**Figure S1**.**10**. J.M. Cruxent site's field notebook of 1969. Drawing (**a**) showing the CX-105 excavation area with the remains of the adult *Notiomastodon platensis*, and the profile from which the carapace (*placa*) fragment and caudal tube (*cola*) of *Glyptotherium* cf. G. *cylindricum* originate. Drawing (**b**) with information regarding the place from which the carapace originates (see **a**). Courtesy Laboratorio de Archeología of the Instituto Venezolano de Investigaciones Científicas.

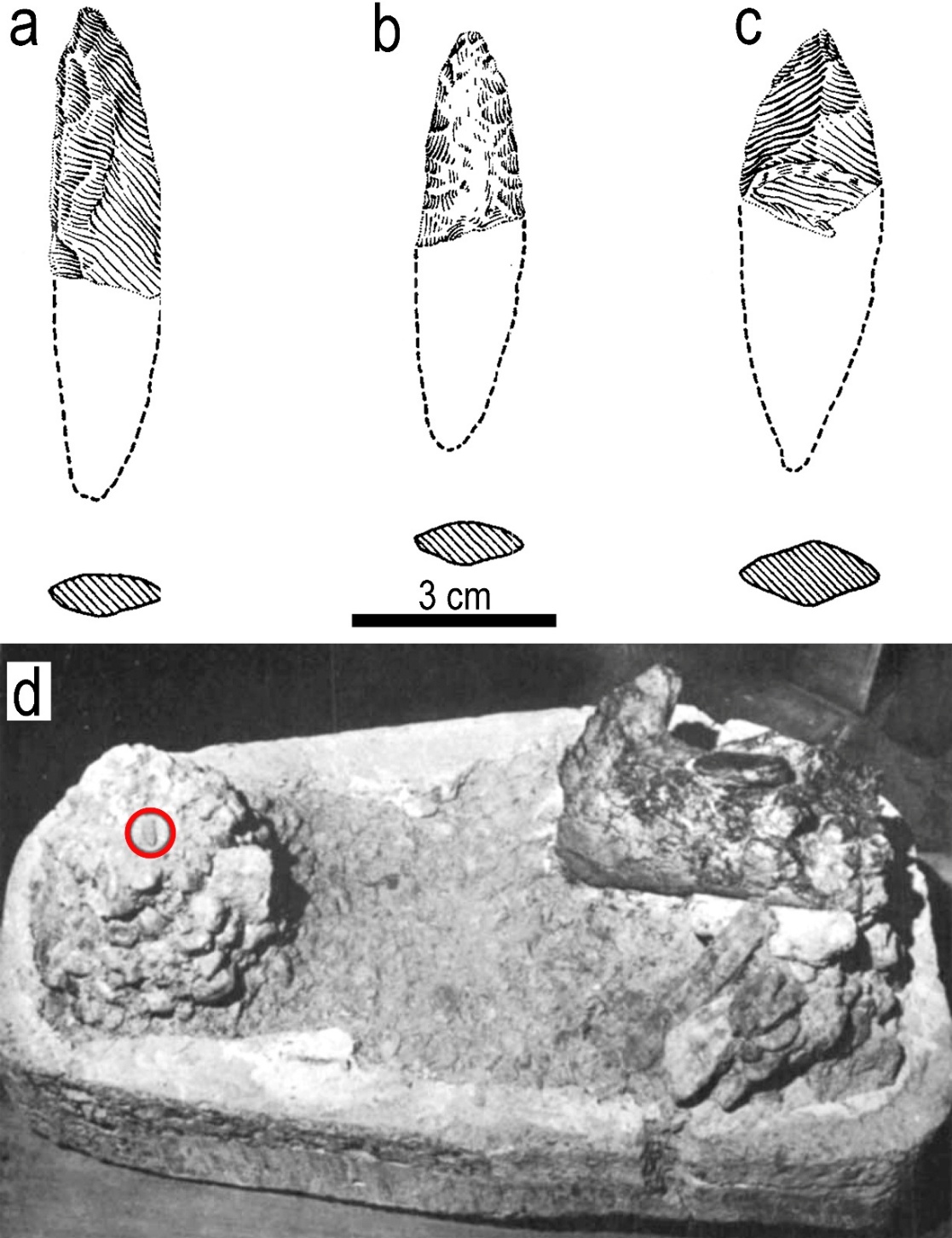

**Figure S1**.**11**. El Jobo-type projectiles found in the 1969 Cucuruchú excavation. Projectiles a and b were lying among the fossil bones. Block of earth (**d**) containing an El Jobo projectile in its original stratigraphic position. Modified images from Cruxent (1970). Projectile in the earth block (**d**) likely corresponds to projectile **a.**

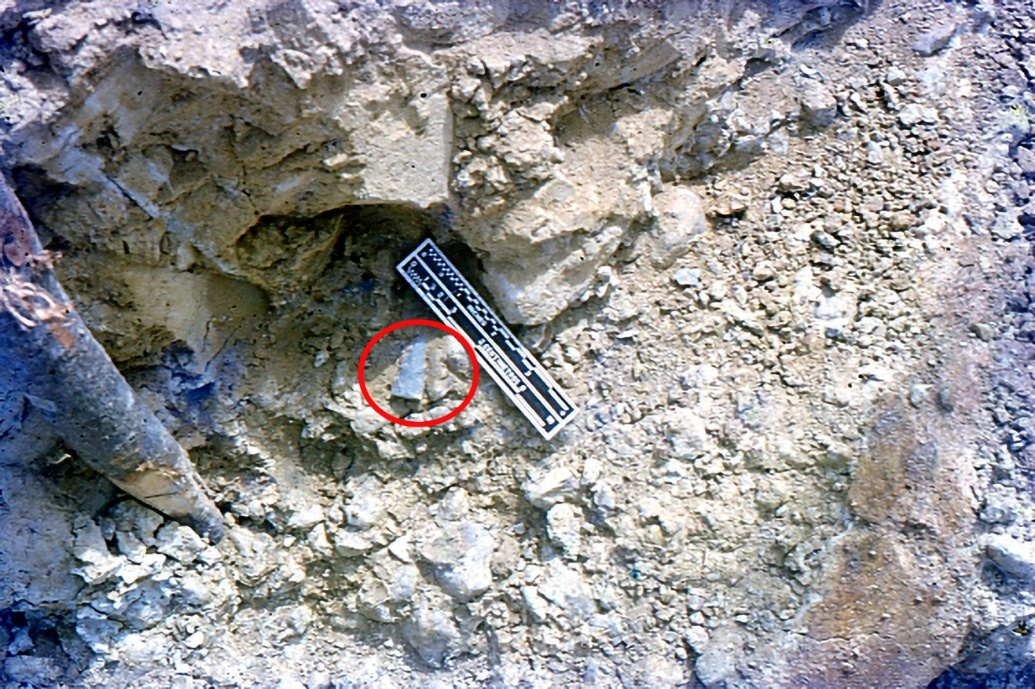

**Figure S1**.**12**. El Jobo-type projectile found in the 1969 Cucuruchú excavation. Image from Fundación José María Cruxent in the custody of the Laboratorio de Archeología of the Instituto Venezolano de Investigaciones Científicas.

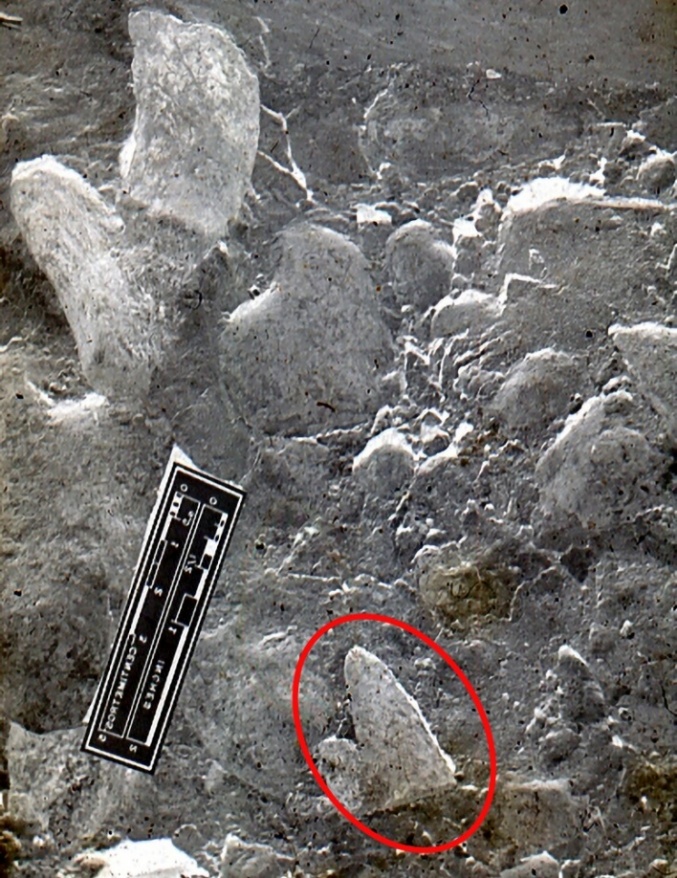

**Figure S1**.**13**. El Jobo-type projectile found among fossils during the 1969 Cucuruchú excavation. Image from Fundación José María Cruxent in the custody of the Laboratorio de Archeología of the Instituto Venezolano de Investigaciones Científicas.

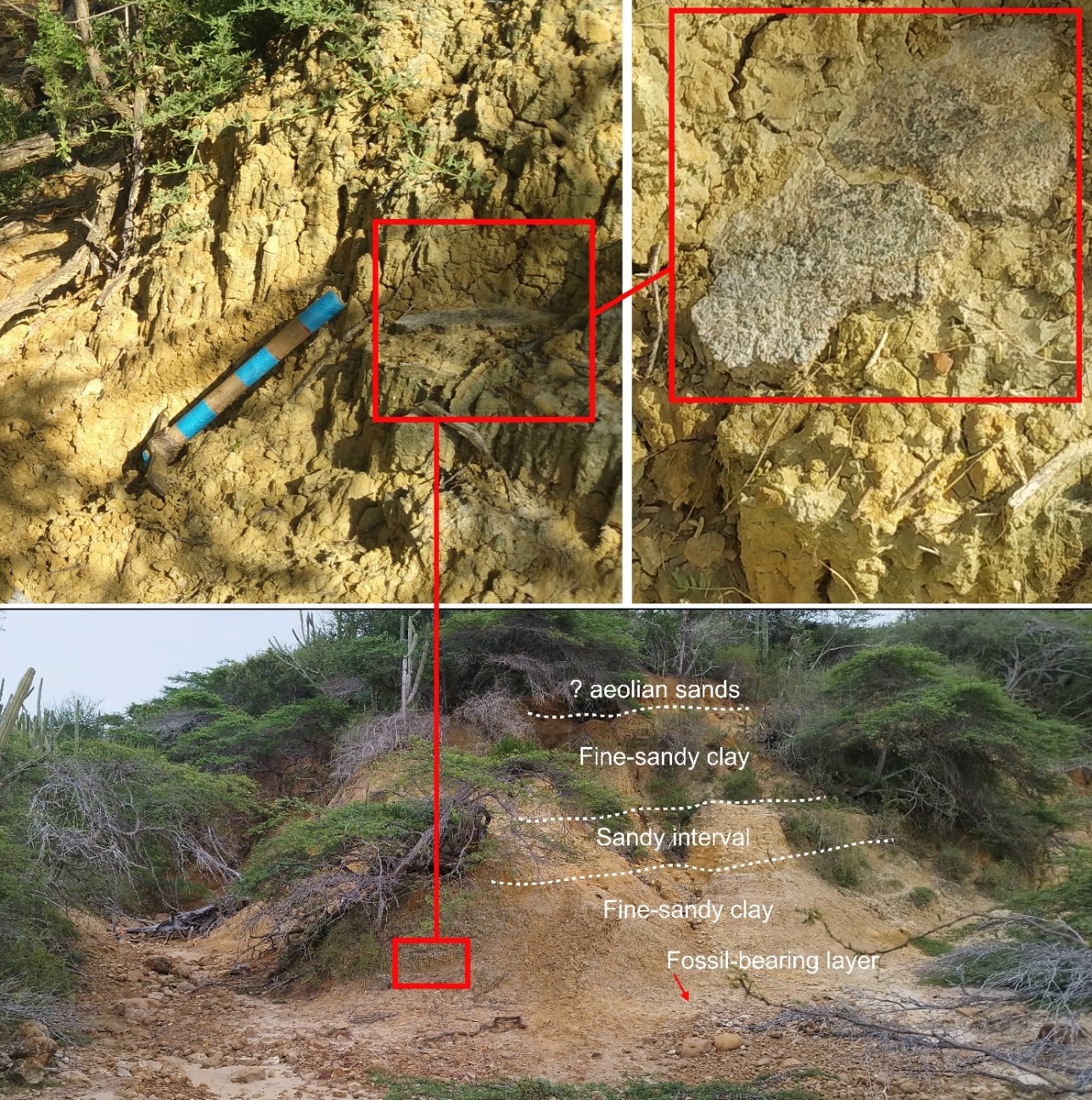

**Figure S1**.**14**. Overview of the Cucuruchú site showing the relative position of the layers. The red rectangles highlight the location where the fragment of the armored carapace MCH-Pv-870 (collected by our team), originating from the base of the fine-sandy clay overlying the fossil-bearing layer.

**Notes on taxonomic assessment**

**XENARTHRA** Cope, 1889

Phyllophaga Owen, 1842

†Megatheriidae Gray, 1821

†*Eremotherium* Spillmann, 1948

†*Eremotherium* *laurillardi* (*Lund, 1842*)

Figure S1.15

*Specimen*s

Remnants of the carcass of a probable individual excavated in 1969 (Figure S1.2). The only elements of this carcass collected in Cucuruchú and found so far in the IVIC collection include a left tibia (IVIC-AP-023) and presumably the astragalus (IVIC- s/n) seen in Figure S1.15.

*Remarks*

The left tibia IVIC-AP-023 (Figure S1.15a–d) has a total length of approximately 590 mm, being somewhat fragmented in the posterior proximal portion, lacking the fibular attachment. Much of the tibia, including exposed cancellous bone, is covered by a black layer and some areas oxidized in ochre, suggesting that this specimen was exposed to high temperatures and fire after its extraction from the site. This could suggest that IVIC-AP-023 may have survived the 1978 fire at the IVIC Archaeology Laboratory, where many artifacts and specimens were damaged. In reference to the astragalus IVIC- s/n it has an approximate anteroposterior length of 245 mm (Figure S1.15e–i). We have not found exact information in the Cruxent site's field notebook regarding the grid from which these remains originated. The Figure S1.2 shows the *E. laurillardi* remains during the extraction process, and they appear to be on the slope, which has been cleaned but not gridded at the time of the picture.

In association with the tibia and astragalus, other postcranial remains can be observed, including at least three vertebrae, some bones difficult to identify due to their perspective in the photograph, and numerous rib fragments. There is no clear information about whether any cranial material was found associated with these remains. Due to the lack of diagnostic characteristics in the Cucuruchú specimens, we tentatively suggest assigning them to *E*. *laurillardi* This assignment is based on the morphology of the tibia and astragalus, like that of other *E*. *laurillardi* individuals collected from Late Pleistocene deposits in Falcón (e.g., Muaco and Taima-Taima sites) and other regions of Venezuela (see Schaub 1935; Aguilera 2006; Carrillo-Briceño 2015; Carrillo-Briceño *et al*. 2016, 2024; Chávez-Aponte 2022, and references therein). Originally Cruxent (1970) assigned these remains to *Eremotherium rusconii*, however the validity of this taxon is questionable, with *E*. *laurillardi* being recognized as the only species of the genus known from the Late Pleistocene of South America (Faure *et al*. 2014; Cartelle *et al*. 2015).

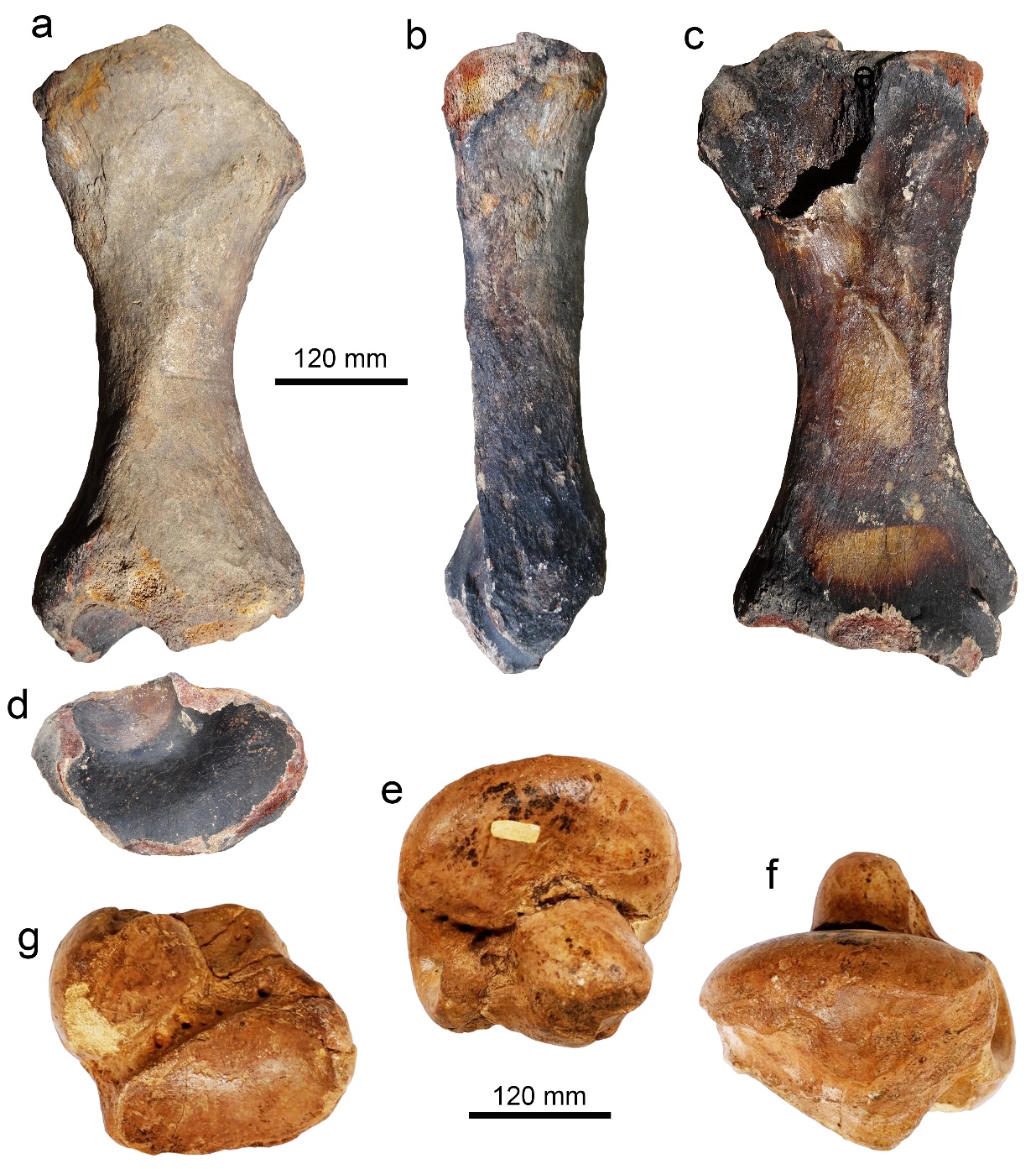

**Figure S1**.**15**. Fossil remains of *Eremotherium* *laurillardi* from Cucuruchú. Left tibia (**a**–**d**: IVIC-AP-023) with burned surfaces probably because of the 1978 fire at IVIC. Right astragalus from Cucuruchú (**e**–**g**: IVIC-s/n). Tibial views: anterior (**a**), lateral (**b**), posterior (**c**), distal (**d**); astragalus views: tibial (**e**), lateral (**f**), calcaneal (**g**).

**Cingulata** Illiger, 1811

†Glyptodontidae Gray, 1869

†*Glyptotherium* Osborn, 1903

†*Glyptotherium cylindricum* (Brown 1912) sensu Gillette and Ray, 1981

*Glyptotherium* cf. *G. cylindricum*

Figures S1.16 and 17

*Specimens*

Glyptodontid remains are the most abundant collected to date at the Cucuruchú site, and these include cranial and postcranial elements. Based on the Cruxent site's field notebook and historical photographs from the 1969 excavation, we believe that the specimens collected in the former excavation correspond to a left femur, a carapace fragment, and a tail fragment of a glyptodontid. The new specimens reported here and collected in the last 10 years include an isolated molar (MTT-V-297), a carapace fragment with 7 articulated osteoderms (MCH-Pv-870), and 31 disarticulated osteoderms (MCH-Pv-8, -876, -877–879, -881, -882, -884, -887–889, -890, -892, -894, -896, -900, -902, -903, -905, -906, -912–914, -916, -917 and MTT-V-172, -339, -341, -444, -727, -916).

*Remarks*

We have not obtained photographs and descriptions of the glyptodont materials collected during the 1969 excavation at Cucuruchú. Cruxent refers to and sketches in his site's field notebook (Figures S1.10) what he calls a glyptodont *placa* (carapace fragment) and originating from what Cruxent describes as an *arcilla* (clay) layer (Figure S1.10a). A few centimeters below the carapace fragment, a fragment of a glyptodont tail was also collected (Figure S1.10a). Both specimens apparently come from a small (unnumbered) excavated section between areas CX-104, CX-105, and CX-108 (Figure S1.7). In the IVIC collection, we have found a fragment of a carapace (IVIC-s/n) and tail of a glyptodont (IVIC-s/n) that could belong to the specimens mentioned in the Cucuruchú site's field notebook (Figures S1.4 and 10). The carapace fragment IVIC-s/n (Figure S1.16a, b) probably belongs to the dorsolateral region and preserves osteoderms characterized by an ornamentation of the external surface consisting of a polygonal central figure surrounded by a groove and a row of polygonal peripheral figures (up to eight in some specimens), the latter always being smaller than the central figure. The central groove has 2 to 4 perforations at the intersection of radial and central grooves; the external surface of the osteoderms is rough. Regarding the tail fragment IVIC-s/n (Figure S1.16c–e), this corresponds to the distal end of the caudal tube with 5 rows of complete osteoderms, which envelop the last vertebrae that does not appear to be fused to the osteoderms, only joined by connective tissue. The first row of the tube was probably in contact with the most posterior caudal ring. Two sets of two osteoderms with dorsal spikes-like (in the second and fourth rows). The distal section of the tube ends in two blunt osteoderms. This specimen was previously mentioned by Aguilera (2006, pp. 42) as coming from Taima-Taima. In excavation area CX-105, between sections -4 and -7, and very close to the remains of the adult *Notiomastodon platensis*, a left femur of a glyptodont is observed (Figure S1.6d).

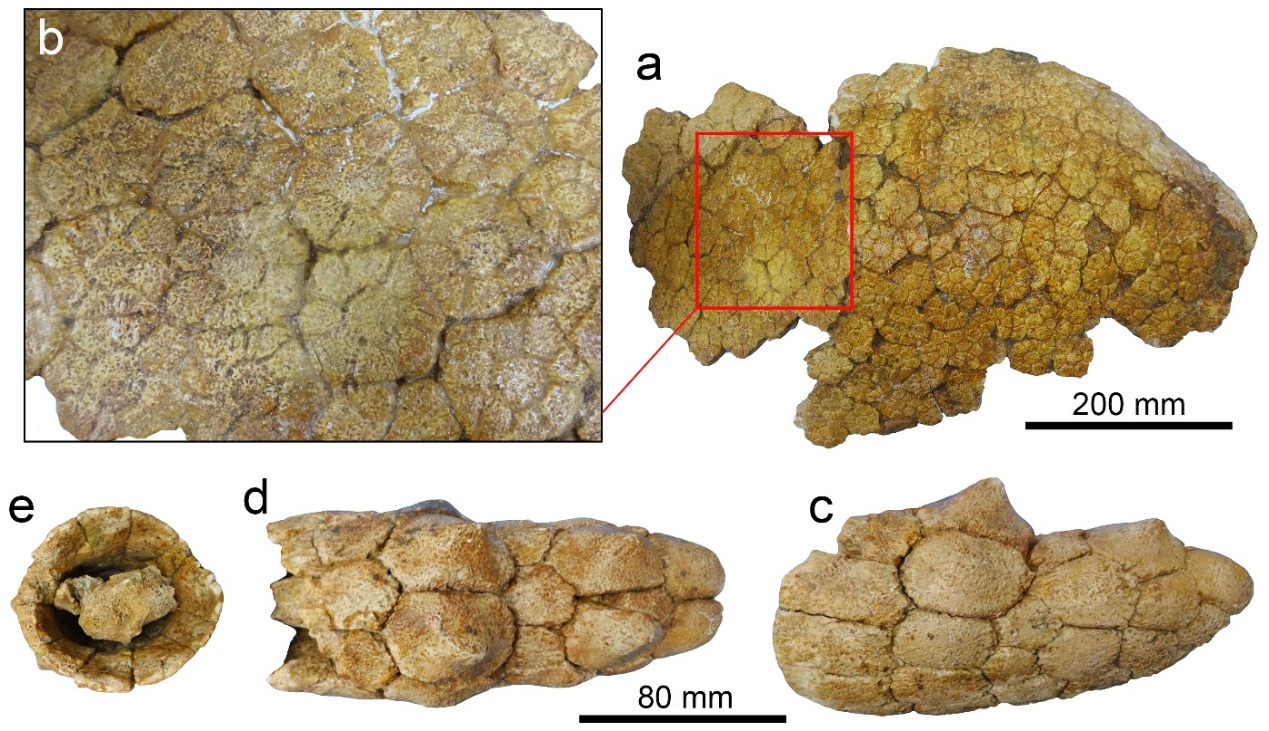

**Figure S1**.**16**. Fragment of the carapace (**a**, **b:** IVIC-s/n) and caudal tube (**c**–**e:** IVIC-s/n) of a *Glyptotherium* cf. *G*. *cylindricum*. These specimens could be those collected at the Cucuruchú site during the 1969 excavation and referred to in Cruxent site's field notebook (see Figs. S1. 10). Views: external (**a**, **b**), left lateral (**c**), dorsal (**d**), section proximal (**e**).

The carapace fragment MCH-Pv-870 with seven articulated osteoderms (Figure S1.17c) and the remaining osteoderms collected by our team (e.g., Figure S1.17) mostly appear to have been affected by erosion and weathering processes due to their exposure to the surface. These osteoderms exhibit morphology like those of the shell fragment mentioned above (Figure S1.16), and some of them also appear to belong to the dorsal and dorsolateral sections of the carapace. Other osteoderms appear to belong to the lateral section (Figure S1.17d, h, n, r, t, v, w), probably the ventral region, the cephalic shield, the carapace margin, and the tail rings. Some isolated osteoderms can reach a length of slightly more than 50 mm. The only cranial element found so far is a relatively well-preserved upper left molar MTT-V-297 (Figure 17a, b) with the typical occlusal pattern that characterizes other specimens of the genus *Glyptotherium* described from Muaco and Taima-Taima (e.g., skull MCNC-Pal-1840; Carlini *et al*. 2008, fig. 2c; 2022, fig. 4). It is not possible to determine whether all the specimens mentioned above are associated and whether they belong to a single individual or to several individuals due to the absence of a detailed stratigraphic context.

In the collection of CIAAP (Universidad Experimental Francisco de Miranda, Coro), there is a specimen (CIAAP-1540) from the tail of *Glyptotherium* cf. *G*. *cylindricum* preserving articulated osteoderms of two partial caudal rings, with three rows of osteoderm each and which was illustrated by Carlini *et al*. (2008, fig. 3e, f) without specifying its provenance. According to the CIAAP catalog, this specimen comes from the Cucuruchú locality and lacks any other information such as collector or year of collection. We do not rule out that this specimen was collected at the Cucuruchú site, outcropping at the surface, sometime after the 1969 excavation.

The remains known from the Late Pleistocene of Falcón state are practically all attributed to the genus *Glyptotherium*, in accordance with a recent revision (Carrillo-Briceño 2015; Zurita *et al*. 2018; Carlini *et al*. 2022). Some specimens, including cranial and postcranial remains and armor elements (e.g carapace, caudal, cephalic shield), have been described for the Muaco and Taima-Taima sites as *Glyptotherium* cf. *G*. *cylindricum* (Carlini *et al*. 2022). The morphology of the osteoderms mentioned here for the Cucuruchú site resembles those described for the Muaco and Taima-Taima sites (Carlini *et al*. 2018). Therefore, the Cucuruchú specimens are referred to *Glyptotherium* cf. *G*. *cylindricum*.

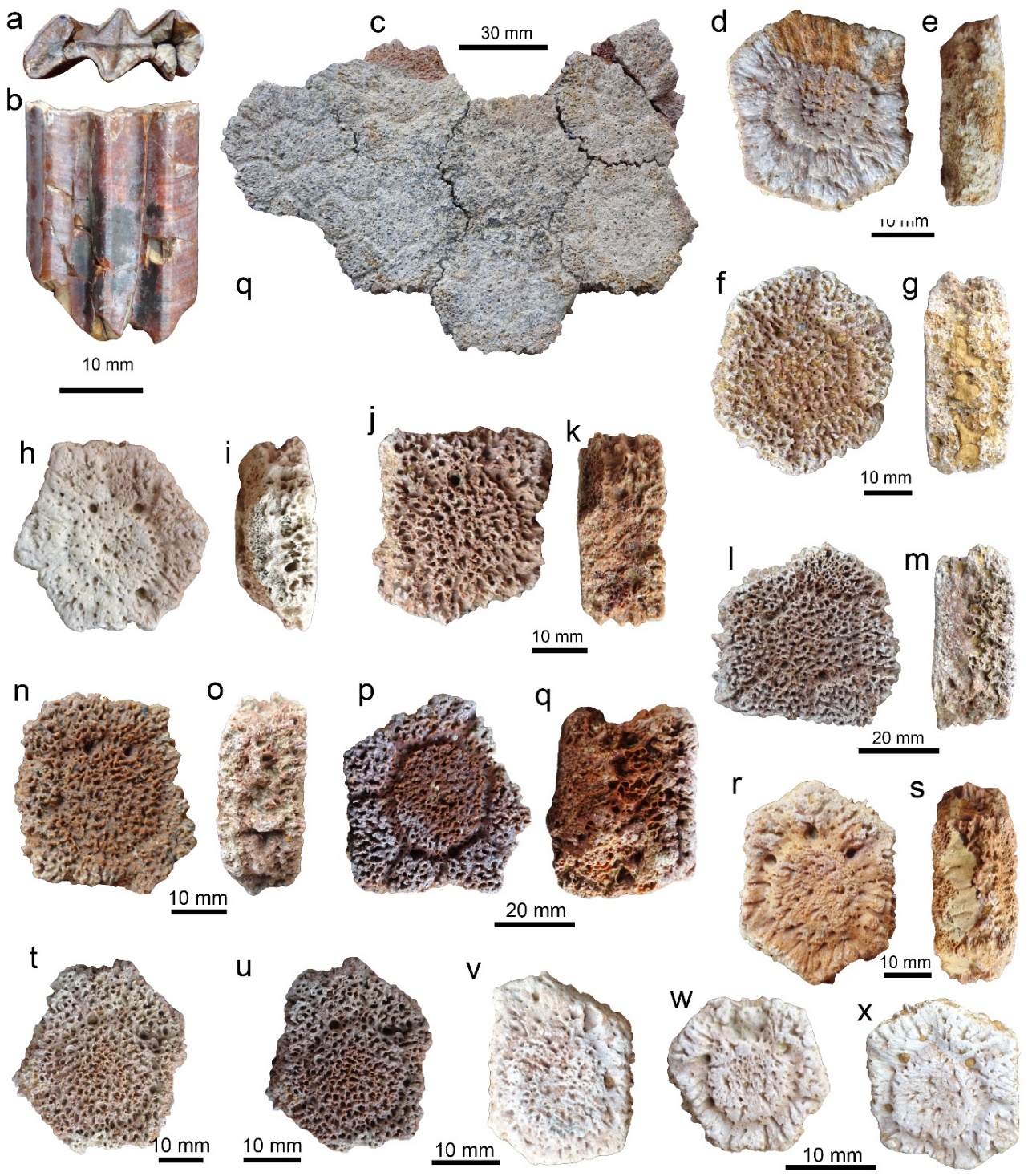

**Figure S1**.**17**. *Glyptotherium* cf. *G*. *cylindricum* remains from Cucuruchú site. Upper left molar (**a**, **b**: MTT-V-297). Fragment of armoured carapace (**c**: MCH-Pv-870). Isolated carapace osteoderms (**d**, **e**: MCH-Pv-912; **f**, **g**: MCH-Pv-903; **h**, **i**: MTT-V-339; **j**, **K**: MCH-Pv-877; **l**, **m**: MCH-Pv-890; **n**, **o**: MCH-Pv-877; **p**, **q**: MCH-Pv-902; **r**, **s**: MCH-Pv-916; **t**: MTT-V-s/n; **u**: MCH-Pv-902; **v**: MCH-Pv-906; **w**: MCH-Pv-913; **x**: MCH-Pv-914). Views: external (**c**, **d,** **f**, **h**, **j**, **l**, **n**, **p**, **r**, **t**–**x**), labial (f), profile (**e**, **g, i**, **k**, **m,** **o**, **q**, **s**), occlusal (**a**).

**†LITOPTERNA** Ameghino, 1889

†Macraucheniidae Gill, 1872

†*Xenorhinotherium* Cartelle and Lessa, 1988

†*Xenorhinotherium bahiense* Cartelle and Lessa, 1988

cf. *Xenorhinotherium bahiense*

Figure S1.18a–i

*Specimen*s

Three mandibular fragments including a dorsoposterior portion, a posterior fragment of the mandibular body, an anterior fragment with a premolar, and an isolated premolar, all with the number MCH-Pv-867.

*Remarks*

The dorsoposterior portion of a left hemimandible preserves the dorsal portion of the ramus with the coronoid and the condylar processes and part of the masseteric fossa (Figure S1.18a, b). The posterior fragment of the mandibular body also belongs to a left hemimandible (Figure S1.18c, d), and preserves the section located between the last molar and the base of the coronoid process. The fragment of the left hemimandible preserves part of the alveolar process with a p3 and three anterior alveolar cavities (Figure S1.18e–g); a well-preserved mental foramen is located at the level of the p3. The p3 has a crown length of about 27 mm. The isolated molar (Figure S1.18h, i) likely corresponds to a left p4; however, this is incomplete on the lingual side. All these fragments come from the fossil-bearing layer and were collected as they outcropped at the surface, very close to each other, having been exposed by erosion. Although the fragments described above exhibit different preservation characteristics (e.g., coloration), we cannot rule out the possibility that they belong to the same individual. The mandibular and dental elements from Cucuruchú (MCH-Pv-867) are here tentatively assigned to cf. *Xenorhinotherium bahiense*, the only macraucheniid so far identified (based on cranial and dental elements) for the Late Pleistocene sites of Muaco and Taima-Taima (Aguilera 2006; Carrillo-Briceño 2015; Carrillo-Briceño *et al*. 2026). The fossil record of *X*. *bahiense* suggests that this was the only macrauchenid species that inhabited northern South America at the end of the Pleistocene (Scherer *et al*. 2009), although disagreements exist over the validity of this taxon, and some authors consider it synonymous with Macrauchenia patachonica (Guerin and Faure 2004).

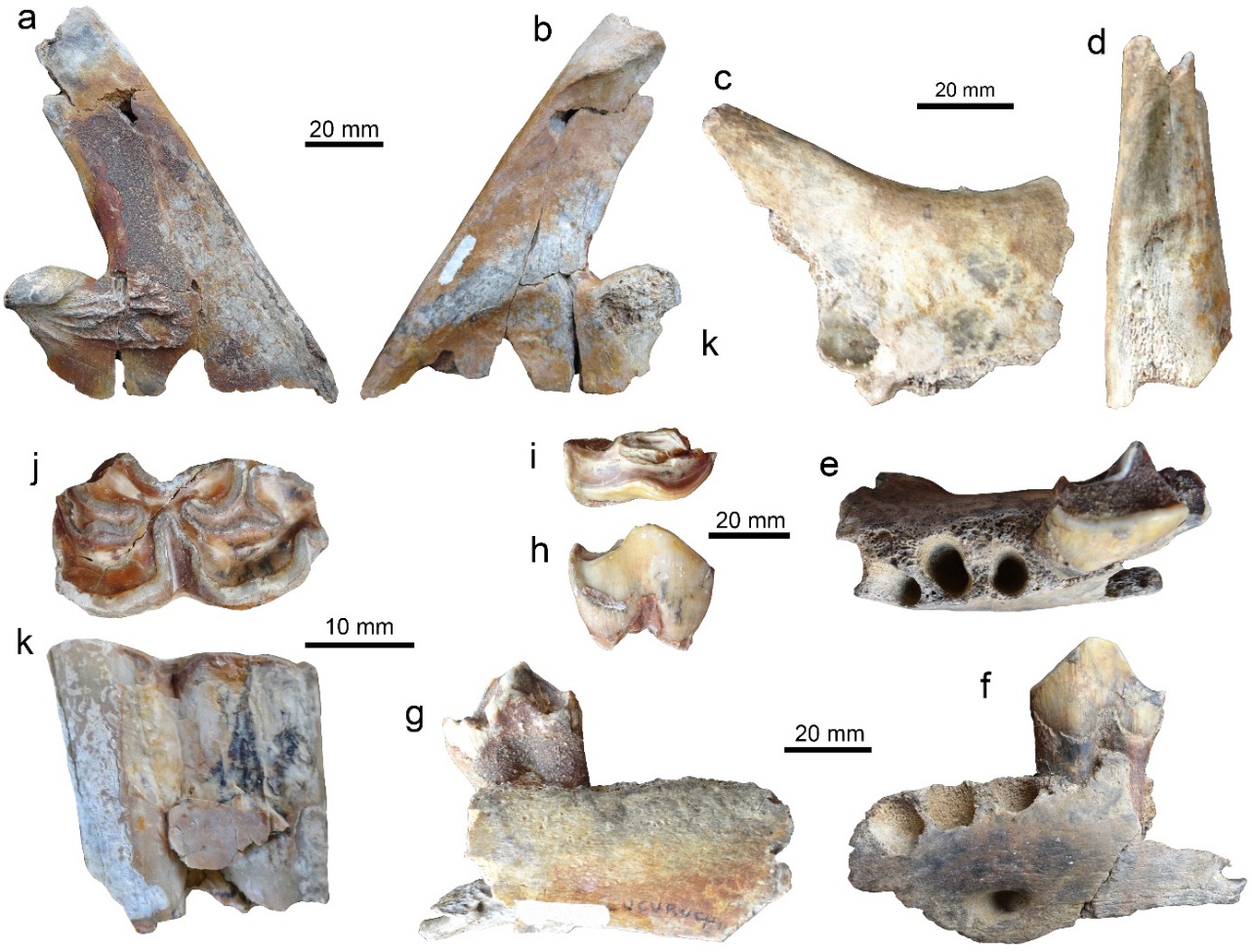

**Figure S1**.**18**. Remains macrauchenids and equids from Cucuruchú site. Fragments of a hemimandible of cf. *Xenorhinotherium bahiense*: dorsoposterior portion (**a**, **b**), posterior fragment of the mandibular body (**c**, **d**), anterior fragment with p3, and isolated ?p4 (all specimens with number MCH-Pv-867). Isolated lower left premolar p3 or p4 (**j**, **k**) of *Equus* sp (MTT-V-323). Views: dorsal (**d**, **e**), labial (?**h**), lingual (**k**), left lateral (**b**, **f**), mesial (**a**, **c**, **g**), occlusal (**i, j**).

**PERISSODACTYLA** Owen, 1848

Equidae Gray, 1821

*Equus* Linnaeus, 1758

*Equus* sp.

Figure S1.18j, k

*Specimen*

Two isolated molars, MCH-PV-728 and MTT-V-323.

*Remarks*

These specimens were collected on the surface of the eroded fossil-bearing layer. MCH-PV-728 corresponds to a molar fragment of determined position and without preserving its occlusal section; radiocarbon dating based on tooth enamel was performed on this specimen. MTT-V-323 is the best preserved (Figure S1.18j, k), especially on the occlusal surface and it probably corresponds to a lower left premolar p3 or p4. The occlusal surface in MTT-V-323 is relatively well preserved with a length of 25 mm and it is characterized by a metaconid and metastylid forming a more rounded double knot, subrectangular to oval protoconid and oval hypoconid, and a triangular and elongated ectoflexid. The presence of a subrectangular to oval protoconid, an oval hypoconid can likely be associated more with *Equus* than *Hippidion* (the other extinct south American genus; see Prado and Alberdi 2017). Since our specimen is not complete and the lack of more diagnostic characters, it is appropriate to keep this assignment tentatively as *Equus* sp.

For the Pleistocene of Falcón State, equid remains assigned to *Equus neogeus* and *Equus santaeelenae*, have been reported for the sites of Muaco, Taima-Taima and Quebrada Ocando (Aguilera 2006; Rincón *et al*. 2006). Other reports of fossil equids from Venezuela have also been referred to by Rincón et al. (2006) and Carrillo-Briceño (2015).

**PROBOSCIDEA** Illiger, 1811

†Gomphotheriidae Hay, 1922

†*Notiomastodon* Cabrera, 1929

†*Notiomastodon platensis* (Ameghino 1888)

Figure S1.19

*Specimen*

Remnants of two individuals excavated in 1969 (Figures. S1.5, 6 and 12a). The only fossils found so far in the IVIC collection and certainly associated with the Cucuruchú site include a complete mandible (IVIC-s/n-CX-108), and two tusks (IVIC-s/n-CX-105).

*Remarks*

The specimens collected in 1969 from the excavated area CX-105 likely belong to the same individual (Figures S1.6). The photographic record shows the mandible with what appears to be both m3, the disintegrated maxilla with upper molars (at least two of them appear to be both M3), and the two tusks. These tusks, deposited at IVIC, still rest on the original sediments and part of the plaster casts used to remove them from the site. The current state of preservation of these tusks shows a level of deterioration and partial fragmentation and disintegration. Despite this, we were able to measure their total length; the longest is approximately 164 cm and the other 134 cm. The longer tusk probably corresponds to the more complete tusk observed in Figure S1.6, while the other was already fragmented during the 1969 excavation. The state of wear of both m3 of the mandible (Figures S1.6d) suggests that this was an adult individual at the time of death. With reference to the complete mandible (Figures S1.5 and 19), this comes from excavated area CX-108 and preserves both m1 and m2 molars. The level of wear of the molars, especially the m1, suggests that this may belong to a subadult or adult individual.

Two Proboscidea species, *Notiomastodon platensis* and *Cuvieronius hyodon*, occurred in South America during the Late Pleistocene–Early Holocene (Alberdi and Prado 2022; Mothé et al. 2010, 2017a); most of their diagnostic features come from dental specimens, including upper tusks and last molars, as well as lower jaw symphysis and tusks (Mothé *et al*. 2016, 2017b). *Cuvieronius* has elongated, twisted, and slightly upcurved upper tusks with a longitudinal enamel band, while *Notiomastodon*'s upper tusks have a great variation in length, robustness, shape, and enamel presence, and are never twisted (Mothé and Avilla 2015). The teeth of both proboscideans are bunodont and quite similar in morphology, differing only in the complexity of the last molars (number of main and accessory cusps), in which *Notiomastodon* had a range of 35–82 cusps and *Cuvieronius* 33–60 cusps (Mothé *et al*. 2016, 2017a, b). Considering these morphological traits, the Cucuruchú specimens present features that fit the diagnostic characteristics of *N*. *platensis*. Some examples include the linear and untwisted tusks with an upcurved shape, and the complexity of some of the molars (e.g., Figure S1.6). *Notiomastodon platensis* has also been reported in the adjacent localities of Muaco and Taima-Taima (Carrillo-Briceño 2015; Carrillo-Briceño *et al*. 2026), and in other Late Pleistocene sites in Falcón (Carrillo-Briceño *et al*. 2024) and Venezuela (Carrillo-Briceño *et al*. 2008; Carrillo-Briceño 2012). Evidence of exploitation in several individuals of *N*. *Platensis* has been reported for Taima-Taima (Bryan *et al*. 1978; Carrillo-Briceño *et al*. 2026).

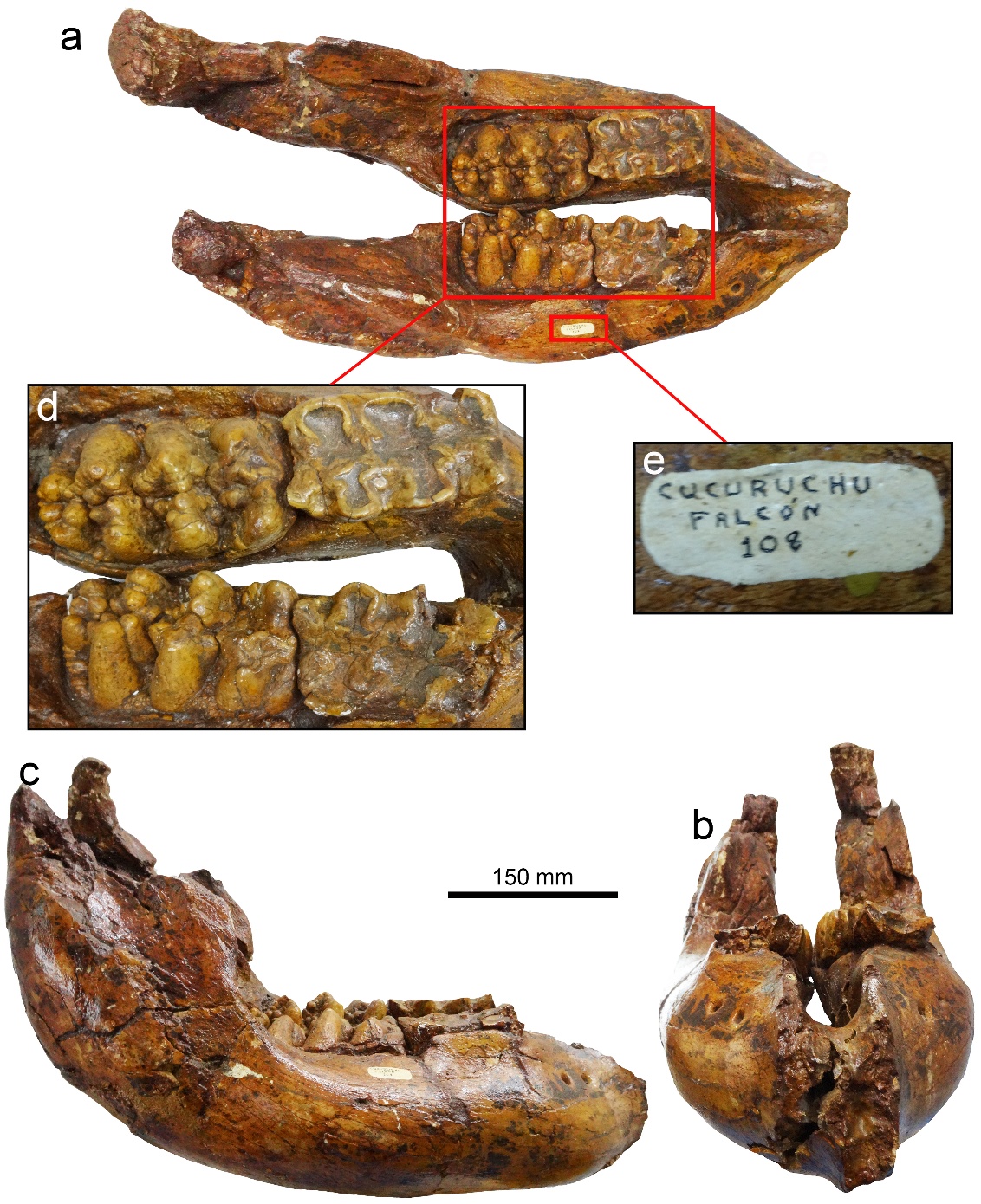

**Figure S1**.**19**. Mandible IVIC-s/n-CX-108 of a probable subadult/adult of *Notiomastodon platensis* coming from the excavated area CX-108. Views: anterior (**b**), dorsal (**a**), right lateral (**c**), occlusal (**d**).

**TESTUDINES** Batsch, 1788

Kinosternidae Agassiz, 1857

*Kinosternon* Spix, 1824

*Kinosternon* sp.

Figure S1.20

*Specimen*s

Two isolated peripheral bones (MTT-V-348 and MTT-V-598).

*Remarks*

MTT-V-348 corresponds to a nearly complete peripheral bone from the anterior margin of the carapace, possibly the right peripheral 3 (Figure S1.20a, b). It measures 15 mm in width and 17 mm in length. MTT-V-598 is a partially preserved peripheral from the bridge region of the shell (Figure S1.20c, d). As preserved, it measures 16 mm in width and 19 mm in length. Both peripherals exhibit small size, a densely microvermiculated bone surface, deep sulci between the marginal scutes, and a pronounced step between the ventral and visceral surfaces. All these features are consistent with the pattern exhibited by species of the genus *Kinosternon*. However, any further taxonomic attribution beyond the genus level is not possible; therefore, we assign the material to *Kinosternon* sp. The occurrence of *Kinosternon* turtles at the Cucuruchú site expands the geographical distribution of the genus during the Late Pleistocene, which was previously known from Mene de Inciarte tar pits, Sierra de Perijá, western Venezuela (Lisett Mesa 2005), and the Pubenza site, Colombia (Cadena *et al*. 2007; Alfonso-Rojas *et al*. 2021).

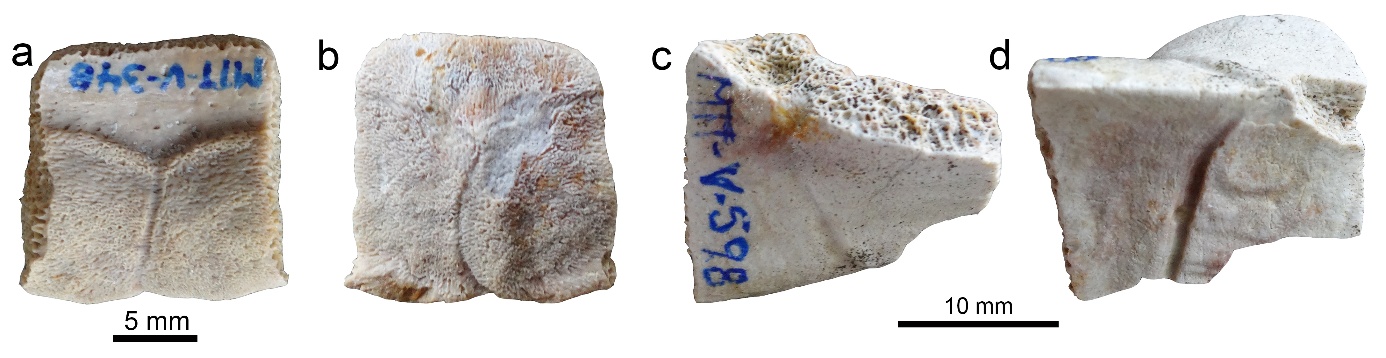

**Figure S1**.**20**. Isolated peripheral bones (**a**, **b**: MTT-V-348 and **c**, **d**: MTT-V-598) of *Kinosternon* from the Cucuruchú site. Views: dorsal (**b**), lateral (**d**), ventral (**c**), ventrovisceral (**a**).
